# Nuclear Speckles Regulate Splicing During Muscle Stem Cell Activation and Aging

**DOI:** 10.64898/2026.05.04.722644

**Authors:** Steve D. Guzman, Pamela Duran, Yu Xiao, Yuxuan Song, Joshua D. Welch, Chuan He, Carlos A. Aguilar

## Abstract

Skeletal muscle contains a population of adult stem cells called satellite cells or muscle stem cells (MuSCs) that are responsible for regeneration after injury. MuSCs utilize gene expression programs to maintain quiescence and differentiate after injury and a key regulator of gene expression is splicing, which uniquely changes when transcripts interact with nuclear speckles (NS). NS are membrane-less biomolecular condensates that phase separate proteins, RNAs and chromatin, but how these organelles regulate molecular processes in MuSCs remains unknown. Herein, we build a comprehensive and systems-level understanding of NS influence on alternative splicing, transcriptional regulation and stem cell function before and after injury and in aging. We establish that NS increased in size and number in MuSCs following injury and influence MuSC activation dynamics. We generated a catalog of isoform-resolved splicing events and linked how RNA interactions with NS amplify splicing completion during the injury response. In old age, MuSCs lose NS, yet shifted towards longer, more completely spliced isoforms enriched for RNA binding protein motifs and multivalency. Our studies unveil evidence that RNA interactions with NS shape stem cell state and regenerative responses but are attenuated in old age.

**HIGHLIGHTS:**

- 3D super-resolution imaging of nuclear speckles in muscle stem cells before and after muscle injury shows intricate relationship with activation
- Isoform-resolved profiling of muscle stem cells shows increases in gene expression and splicing during injury response
- Mapping RNA interactions with nuclear speckles shows RNAs undergo strongest splicing when proximal to nuclear speckles
- Old aged muscle stem cells lose nuclear speckles and display aberrant splicing, with longer transcripts, more exons, and increased RNA binding protein motifs

## Main

Muscle stem cells (MuSCs) are tissue resident stem cells that maintain quiescence^1^ and differentiate to regenerate tissue after injury^2^ or self-renew^3^. The specific activation and/or repression of genes utilized by MuSCs to execute regenerative functions are critically guided by the processing of RNAs. RNA processing^4^ is critical for MuSC processes^5,6^ which is in part regulated by nuclear speckles (NS)^7^. NS are membrane-less biomolecular condensates that phase separate chromatin, RNAs^8,9^ and proteins^10^ and are strongly associated with pre-mRNA processing and splicing ^11,12^. Interruptions to RNA homeostasis and splicing contribute to multiple pathologies^13^, but how NS and RNA processing regulate transcriptional diversity in MuSCs remains understudied.

NS contain different types of RNAs, RNA binding proteins and RNA processing factors^14^. Scaffold proteins in NS such as SON^15^ and the serine/arginine repetitive matrix 2 protein (SRRM2) ^16^ self-assemble and promote phase separation into membraneless condensates due to their intrinsically disordered regions. SON and SRRM2 form the “core” of NS and serine/arginine splicing factors and heterogeneous nuclear ribonucleoprotein splicing factors form the “shell” of NS ^17,18^. Other spliceosome proteins^7^ and RNA polymerase II decorate the surface of the nuclear speckle shell and when loci physically interact with NS, splicing reactions are catalyzed and enhanced ^19^ due to compartmentalized reactions^20^. The loss of SRRM2^21^ or SON^15^ drive aberrant nuclear speckle morphology, splicing defects, and reduce chromatin interactions in active compartments^22^ and RNA expression levels. However, no studies have evaluated how RNAs and their interactions with RNA binding proteins in NS^23^ contribute to transcript structure and fate in MuSCs or adult stem cells more broadly, nor how nuclear speckle dynamics are altered during aging.

Herein, we demonstrate how nuclear NS regulate MuSC splicing and gene expression programs before, after injury and in aging. We show NS are required for MuSC proper activation and regeneration, and that old age is associated with smaller and fewer NS. We profile molecular interactions between RNAs and NS before and after injury as well as in aging and show NS shape variations in splicing that contribute to MuSC regenerative potential.

## Results

### Three-dimensional super-resolution imaging of nuclear speckles in muscle stem cells before and during regeneration reveals an association with activation

To begin to understand how NS change in response to MuSC states, we isolated MuSCs from hind limb muscles (gastrocnemius, tibialis anterior, and quadriceps) of young mice (3-5 months) at 0- and 7-days post-injury (dpi) via injections with barium chloride (BaCl_2_) (**Fig. 1A**). We immuno-stained MuSCs for SRRM2 and a nuclear envelope protein (Lamin A/C) (**Fig. 1B)** and performed 3D super-resolution microscopy. We structurally separated each nanoscopic organelle from its neighbors in 3D and found uninjured MuSCs have 10.5% smaller (**Fig. 1C** and **Fig. S1A**) and 52% less (**Fig. 1D**) NS than injured / differentiating MuSCs. These results show MuSC activation leads to increased nuclear speckle number and size.

**Figure 1.**
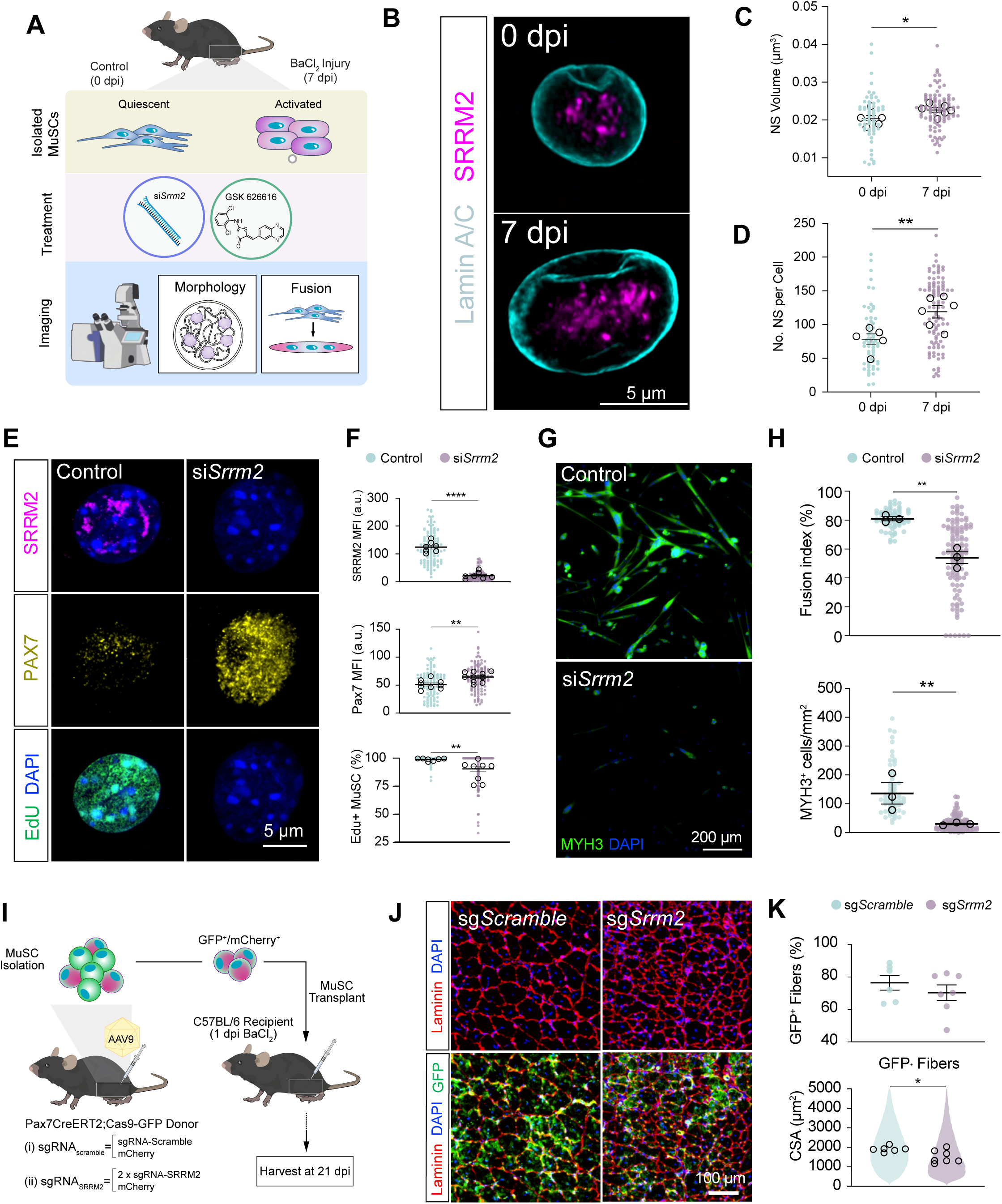
Nuclear speckles support muscle stem cell (MuSC) activation. (A) MuSCs were isolated from young mice across different conditions as described and processed for the assessment of nuclear speckle morphology and the functionality of MuSCs. (B) Representative images of NS at 0- and 7-days post-injury (dpi). Cyan= Lamin A/C, Magenta= SRRM2. C) Quantification of the volume of individual NS. Single cells= 10-30 per replicate (average of individual nuclear speckle volume per cell), replicates= 5-6 animals per group. (D) Quantification of the number of NS across groups (average of nuclear speckle number per cell). Single cells= 10-30 per replicate, replicates= 5-6 animals per group. (E) Single-cell images of SRRM2 (top), PAX7 (middle), and EdU (bottom) in lipofectamine control and *Srrm2* knockdown. (F) Quantification of the total mean fluorescence intensity (MFI) of SRRM2 (top, single cells=12-35 per replicate, replicates=7 wells per group), Pax7 (middle, field of views= 10-18 per replicate, replicates=9-10 wells per group), and the percentage of EdU^+^ MuSCs (bottom, field of views= 10-18 per replicate, replicates= 7-8 wells per group) between lipofectamine control and SRRM2 knockdown. (G) Representative images of differentiated and fused MuSCs between lipofectamine control and *Srrm2* knockdown (Green= Myh3, Blue= DAPI). (H) Quantification of fusion index and density of MYH3^+^ cells (field of views= 20-48 per replicate, replicates= 3 animals per group) between groups. (I) Schematic describing transplantation of SRRM2-KO cells to injured muscle. Pax7CreERT2-Cas9-GFP mice were injected with tamoxifen for 5 days to activate Cas9. Adenoassociated virus 9 (AAV9) either loaded with two gRNAs targeting SRRM2 or a non-targeting/scramble gRNA were injected intramuscularly to the tibialis anterior (TA) and gastrocnemius muscles. Three weeks after delivery, GFP^+^ cells were isolated and injected to a BaCl_2_ injured muscle of wild type mice. (J) Representative images of a cross-section of TA muscle at 21 dpi previously transplanted with sgScramble GFP^+^ MuSCs (top) or SRRM2-KO GFP^+^ MuSCs (bottom) (Red= Laminin, Green= GFP^+^, Blue= DAPI). (K) Quantification of the percentage of GFP^+^ fibers between scramble and SRRM2-KO groups (top). Quantification of the cross-sectional area of GFP^+^ fibers, violin plot represents the overall distribution of all replicates between scramble and SRRM2-KO groups and circles indicate the median of each replicate (bottom, replicates= 6-7 animals per group). *p-* values derived from student’s t-test and Mann-Whitney test. \**p*<0.05, \*\**p*<0.01, \*\*\*\**p*<0.0001; mean ± SEM.

We reasoned that if NS are critical for MuSC activation, then knockdown of *Srrm2* would reduce ability to activate. We knocked down Srrm2 with siRNAs in MuSCs ex vivo (**Figs. 1A** and **Fig. S1B**) and detected that loss of SRRM2 resulted in increases in Pax7, a marker of MuSC quiescence ^24^, and reduction in EdU^+^ incorporation (**Figs. 1E-F**). Differentiation of MuSCs into myotubes and immunostaining for myosin heavy chain 3 (MyH3) further revealed that the loss of *Srrm2* reduced fusion index and the number of MyH3^+^ cells (**Fig. 1G-H**), as well as the number of nuclei per myotube compared to controls (**Fig. S1C**). We then assessed if other nuclear speckle scaffolding proteins (SON) were impacted by the loss of *Srrm2*. In line with prior studies^15,21^ showing that SRRM2 is critical for NS organization and that perturbation of SRRM2 disrupts speckle integrity, we observed a 10% decrease in SON-positive speckle size and a 41% increase in SON-positive speckle number after *Srrm2* knockdown, consistent with speckle fragmentation (Fig. S1D).

To determine if increases in NS size and number potentiate MuSC activation, we isolated and cultured MuSCs in the presence of a small molecule (GSK-626616) that inhibits the dual specificity kinase 3^23^ (DYRK3), which has previously been shown to dissolve NS during interphase^25^ (**Fig. 1A**). MuSCs treated with GSK-626616 increased SRRM2 size and reduced Pax7 intensity when compared to control MuSCs (**Fig. S1E**). Taken together, these results suggest NS may be critical to support MuSC activation.

### Transplantation of muscle stem cells after nuclear speckle modification shows defects in regenerative potential

An enacting measure to assess changes in the regenerative potential of MuSCs due to gene loss is through transplantations^26^. We generated a murine model that facilitates inducible editing of MuSCs in-situ using CRISPR-Cas9 (**Fig. S1F**). This system uses a Cre-specific driver in MuSCs (Pax7^CreERT2^-Rosa26^LSL-Cas9-GFP^ or PROC) to activate Cas9^27^ facilitating knockout of any gene through provision of short guide RNA (sgRNA). We constructed adeno-associated vectors (AAVs) that contain a fluorescent marker (mCherry) to track transfection efficiency, and two sgRNAs to target the SRRM2 locus or non-targeting/scramble sgRNAs as the control. We administered the AAVs through intra-muscular injection to tibialis anterior (3x10^11^ vg) and gastrocnemius (4.5x10^11^ vg) muscles of PROC mice, and three weeks after injection, extracted MuSCs using fluorescence-activated cell sorting (FACS) (**Fig. S1F)**. We observed transfection efficiency (∼20%) that was consistent with previous studies^28,29^ (**Fig. S1H**). To confirm CRISPR gene editing, we isolated DNA from mCherry^+^/GFP^+^ cells and Sanger sequencing demonstrated mixed base calls of approximately 125 bp length beginning at the expected Cas9 cut site for knockout compared to controls (**Fig. S1F and S1G**). The isolated mCherry^+^/GFP^+^ cells also exhibited a 48% reduction in SRRM2 for knockout compared to controls (**Fig. S1H**), demonstrating in situ genome editing of MuSCs and loss of SRRM2. We next delivered AAVs and isolated mCherry^+^ MuSCs from scramble and SRRM2 knockout groups followed by transplantation into pre-injured tibialis anterior muscles^30^. Three weeks after transplantation, injured tissues showed non-significant decreases in the percentage of GFP^+^ and GFP^-^ myofibers and significant decreases in the cross-sectional area of GFP^+^ myofibers for SRRM2 knockout compared to controls (**Figs. 1I-K and Fig. S1I-J**). Combining these results show SRRM2 loss reduces the reparative function of MuSCs.

### Paired short- and long-read single-cell sequencing reveals coordinated changes in splicing completion and isoform usage during MuSC activation

To begin to link NS and splicing to MuSC function, we profiled single cell gene expression from pooled hind-limb muscles of uninjured and 7 days after BaCl_2_ injury using 10x Genomics 3′ chemistry. We sequenced the barcoded cDNA on Illumina NovaSeq X (short read) and Oxford Nanopore PromethION (long read) to generate matched datasets^31^ (**Fig. 2A**). After quality control and filtering^32^, we generated 42,639 cells with paired libraries and a mean of 27,731 short reads / cell and 17,557 long reads / cell across conditions (**Fig. S2A**). Dimensionality reduction^33^ and Louvain clustering followed by cluster annotation by marker genes revealed cell types that were congruent across short-read genes, long-read genes, and long-read isoforms (**Fig. 2B**; **Figs. S2B-C**). Short- and long-read UMIs were highly correlated (r = 0.88, slope = 1.22; **Fig. 2C**), gene detection scaled similarly with UMI depth for both datasets (**Fig. S2D**), and mean expression profiles showed close agreement globally (r = 0.92, slope = 1.01; **Fig. 2D**) and for each cell-type (Pearson correlations typically ≥ 0.87, **Fig. S2E**). Isoforms were called from the long-read data with IsoQuant^34^ and read-length distributions for long-read datasets were >5x longer than the short-read length (857 bp compared to 150 bp, **Fig. S2F**). We re-embedded MuSCs from 0 and 7 dpi using short-read gene expression and used Slingshot^35^ to infer a continuous activation trajectory from quiescent to activated and differentiating states (**Fig. 2E**). We then overlaid long-read isoform expression for *Pax7*, *Myf5*, *Myod1*, and *Myog*, and detected the same trajectory indicating that isoform-level profiles are concordant with the gene expression–based ordering of MuSC states (**Fig. 2F**).

**Figure 2.**
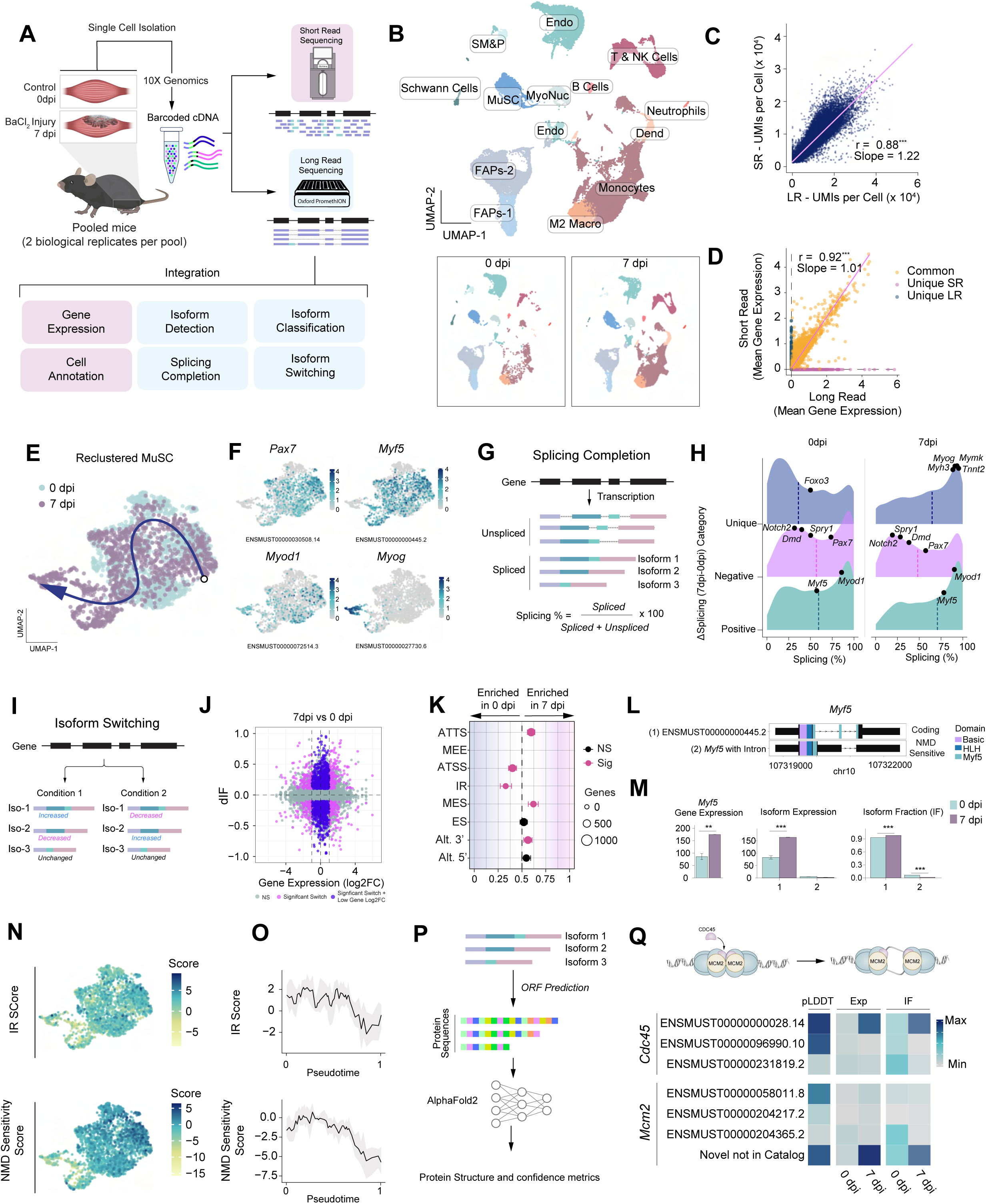
Paired short- and long-read single-cell profiling links splicing completion, isoform switching, and protein structure during MuSC activation. (A) Schematic of experimental design showing isolation of mononuclear cells from control (0 dpi) and BaCl₂-injured (7 dpi) hind-limb muscles, 10x Genomics 3′ capture, splitting of barcoded cDNA for Illumina short-read and Nanopore long-read sequencing, and integrated analyses (gene expression, cell annotation, isoform detection/classification, splicing completion, and isoform switching). (B) UMAP of integrated single-cell data colored by annotated cell type (top), with separate UMAPs colored by injury condition (0 dpi, 7 dpi; bottom). (C) Scatterplot of per-cell UMI counts from short-read versus long-read libraries, with regression line, slope, and Pearson correlation. (D) Scatterplot of mean gene expression measured by long reads versus short reads, with points colored by detection category (common, short-read only, long-read only) and regression line and Pearson correlation. (E) UMAP embedding of the MuSC subset from 0 dpi and 7 dpi colored by condition, with Slingshot-inferred activation trajectories overlaid. (F) UMAP feature plots for long-read isoform expression of *Pax7*, *Myf5*, *Myod1*, and *Myog* across MuSCs. (G) Schematic illustrating the junction-weighted splicing percentage metric, in which fully spliced, partially spliced, and unspliced reads are combined to compute per-gene splicing completion. (H) Density plots of junction-weighted splicing percentage for 0 dpi (left) and 7 dpi (middle) MuSCs and of ΔSplicing% (7 dpi − 0 dpi; right), with example quiescence and activation genes labeled. (I) Schematic representing isoform switching, in which the relative use of specific isoforms can differ between conditions. (J) Scatterplot of Δ isoform fraction (dIF) versus gene-level log₂ fold change (7 dpi versus 0 dpi) for all tested isoforms, with points colored by switch significance and gene-level change category. (K) Dot plot summarizing significant isoform switches grouped by alternative splicing class (ATTS, ATSS, MEE, IR, MES, ES, Alt. 3′, Alt. 5′); x axis indicates enrichment at 0 dpi or 7 dpi, point size reflects the number of genes, and color denotes significance. (L) Genome browser view of Myf5 showing long-read–defined isoforms, retained intron, predicted NMD-sensitive isoform, and annotated protein domains. (M) Bar plots for *Myf5* showing total gene expression, isoform-specific expression, and isoform fractions for Isoform 1 and Isoform 2 at 0 dpi and 7 dpi. (N) UMAP feature plots of MuSCs colored by module scores derived from isoform switches associated with intron retention (IR score), NMD-sensitive isoforms (NMD sensitivity score), and protein-domain gains or losses (protein domain score). (O) Plots of IR (top) and NMD sensitivity (bottom) scores along Slingshot pseudotime, with smoothed trend lines and shaded variability. (P) Schematic of the long-read–based structural workflow: open reading frame prediction for switched isoforms, translation to protein sequences, and AlphaFold2 prediction of structures and per-residue confidence (pLDDT). (Q) Schematic of the CDC45–MCM2 DNA replication pathway (top) and heatmap of representative isoform switches in *Cdc45* and *Mcm2* (bottom), showing relative AlphaFold2 pLDDT, isoform expression, and isoform fraction across 0 dpi and 7 dpi.

The quantification of RNA splicing completion and exon ligation has not been systematically defined as MuSCs transition from quiescence to activation. We quantified splicing completion from the long-read data by calculating a per-gene splicing percentage using a junction-weighted metric that treats partially spliced reads as only partially contributing to the spliced pool (**Fig. 2G**). We first stratified genes that were uniquely expressed at either 0 dpi or 7 dpi. Unique transcripts in quiescent MuSCs (0 dpi) such as *Foxo3* were largely incompletely spliced, whereas unique transcripts at 7 dpi showed a clear enrichment for nearly fully spliced reads and included myogenic differentiation genes such as *Myog*, *Mymk*, *Myh3*, and *Tnnt2* (**Fig. 2H**). We then asked whether activation alters splicing of genes that are present in both states by restricting the analysis to genes detected at both 0 dpi and 7 dpi and examining ΔSplicing% (7 dpi − 0 dpi). Genes with negative ΔSplicing% were enriched for quiescence regulators including Pax7, Spry1, and Notch2, indicating that these transcripts not only decrease in expression but also become less completely spliced upon activation. In contrast, genes with positive ΔSplicing% showed partially spliced distributions at 0 dpi that shifted toward uniformly high splicing at 7 dpi, paralleling the behavior of uniquely expressed differentiation markers (**Fig. 2H**). We next related splicing behavior to changes in mRNA abundance (**Fig. S2H**). Transcripts that were transcriptionally upregulated at 7 dpi exhibited the highest splicing percentages, whereas downregulated, quiescence-associated genes remained incompletely spliced. Together, these data indicate that activation-linked increases in gene expression are accompanied by amplified exon ligation, while the quiescent transcriptional program persists in a more incompletely spliced state.

We next tested whether the relative usage of specific isoforms differs between conditions using IsoformSwitchAnalyzeR^36^ and DEXSeq^37^ and detected 2,027 significant isoform switches, involving 1,782 isoforms across 1,337 genes (**Fig. 2I-J**). We classified genes with switched isoforms by alternative splicing category, and found that intron retention and alternative transcription termination sites were enriched among genes with higher isoform usage at 0 dpi. In contrast, at 7 dpi, genes with switched isoforms were enriched for mutually exclusive exons and alternative 3′ sites (**Fig. 2K**). As an illustrative case, *Myf5* displayed a coding isoform and a novel intron-retaining isoform predicted to be nonsense-mediated-decay (NMD) sensitive based on a premature termination codon within the retained intron (**Fig. 2L**). At 0 dpi, ∼25% of *Myf5* isoforms were NMD-sensitive, while at 7 dpi, <5% of isoforms were NMD-sensitive. The switch in isoform fractions at 7 dpi is consistent with increased gene and isoform-level expression of the Myf5 coding transcript (**Fig. 2M**).

We next asked whether isoform switching events were associated with broader changes in coding potential and transcript quality. Across all switched isoforms, the loss of intron retention was enriched at 7 dpi, whereas NMD-sensitive isoforms were preferentially associated with 0 dpi, while gain or loss of annotated protein domains showed no strong global directional bias (**Fig. S2I**). We constructed module scores from genes harboring intron-retention–associated switches and NMD-sensitive switches and overlay onto the UMAP revealed overlap with uninjured MuSCs and declines along pseudotime. Together, these results indicate that isoform programs favoring intron retention and decay-prone transcripts are most prominent in quiescent cells and are progressively resolved during MuSC activation (**Figs. 2N–O**).

To evaluate the structural consequences of differences in the isoform programs, we integrated AlphaFold2 structure prediction^38^ with our long-read–defined isoforms. For each isoform with a predicted open reading frame, we translated the coding sequence into predicted local distance difference test (pLDDT) scores (**Fig. 2P**). Higher pLDDT suggests greater confidence that an isoform encodes a more ordered, well-defined local fold, whereas lower pLDDT can reflect increased disorder, or reduced confidence in the predicted structure^38^. Globally, isoforms that increased in usage with activation trended toward higher pLDDT values than isoforms that decreased (**Fig. S2J**). For example, we detected two core components of the CMG replicative helicase^39^ (*Cdc45* and *Mcm2*) increased isoform expression and a shift in isoform fraction toward isoforms with higher predicted pLDDT values after injury (**Fig. 2Q**). The assembly of CDC45 with MCM2 is critical for helicase activation and supports our analysis can capture selective usage of isoforms predicted to encode more confidently folded protein products associated with functional DNA replication complexes that are required for activated MuSCs to enter the cell cycle.

### Identification of RNA interactions with nuclear speckles in muscle stem cells before and after injury

To identify RNAs that interact with NS in MuSCs, we applied ARTR-seq^40^, an antibody-guided reverse transcription assay in which a primary antibody against a RNA binding protein recruits reverse transcriptase and biotinylated nucleotides to nearby RNAs, followed by cDNA synthesis, purification and sequencing of biotinylated cDNA (**Fig. 3A**). We used an SRRM2 antibody to label nuclear speckle-proximal transcripts and an IgG antibody as a negative primary control. In situ validation and imaging confirmed biotin signal was strongly enriched at SRRM2 foci relative to the negative primary antibody, and ARTR-seq tracks showed robust enrichment of the nuclear speckle-associated long noncoding RNA Malat1^12,41^ in both 0 dpi and 7 dpi samples (**Fig. S3A–C**). Previous mapping of nuclear speckle associated transcriptomes revealed transcripts that were co-located with NS contained more introns^19^. In line with these results, our ARTR-seq profiles revealed intronic enrichment for transcripts interacting with SRRM2 compared to negative primary IgG controls at both 0 dpi and 7 dpi (**Fig. 3C, Fig. S3D**). We further validated nuclear speckle proximity of transcripts for *Myod1*, which displayed enrichment of introns compared to exonic coverage that was more similar across conditions (**Fig. 3B**). To validate our ARTR-seq data, we performed RNA-scope for *Myod1* together with SRRM2 immunostaining in MuSC nuclei. In line with our ARTR-seq data, *Myod1* RNAs were concentrated near SRRM2-defined NS at both time points (**Fig. 3D-E**). These data confirm we can identify transcripts that are proximal to NS in MuSCs. We next asked which genes show differential nuclear speckle association before and during regeneration. Intron-level ARTR-seq analysis identified nuclear speckle-enriched gene sets that were preferentially associated with either 0 dpi, shared across time points, or enriched at 7 dpi (**Fig. 3C**). Intron-enriched genes at 0 dpi included regulators linked to quiescence and stress signaling such as *Gadd45b*, *Notch1*, and the splicing factor *Srsf10* as well as nuclear speckle and chromatin regulators such as *Srrm2*, *Srsf5*, and *Hdac1*. In contrast, 7 dpi intron-enriched genes included myogenic activation and differentiation factors such as *Myo1g*, *Myh3* together with signaling and cytoskeletal regulators such as *Ddit4*, *Tgfb1i1*, *Tnfrsf14*, and *Ptprv*.

**Figure 3.**
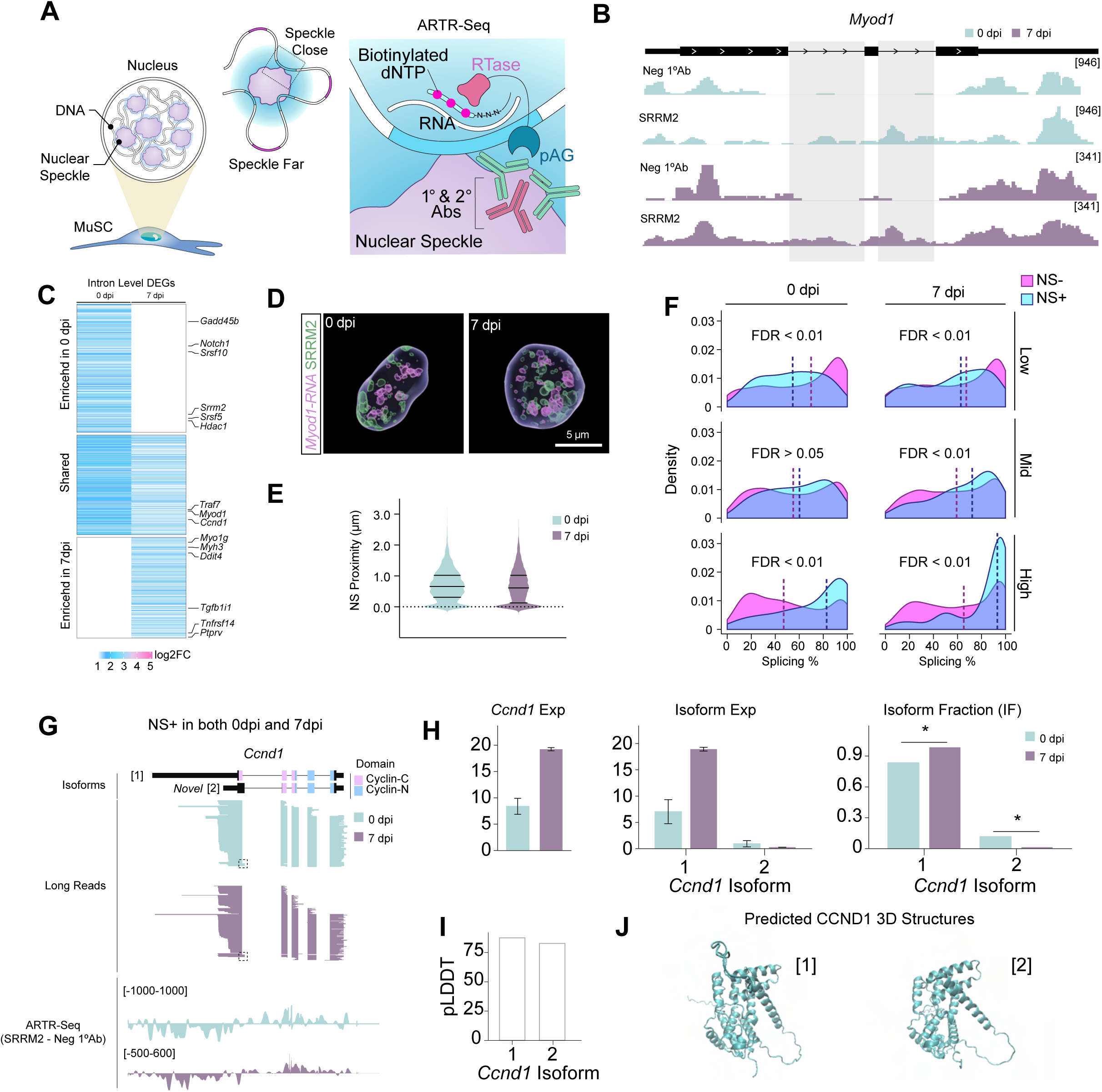
ARTR-seq maps nuclear speckle–associated transcripts and links nuclear speckle proximity to splicing completion. (A) Schematic of ARTR-seq in MuSCs, in which a primary antibody against SRRM2 (or a negative primary antibody control) recruits reverse transcriptase and biotinylated nucleotides to RNAs near NS, followed by purification and sequencing of biotinylated cDNA. (B) Genome browser views of *Myod1* showing gene structure (top) and ARTR-seq tracks for negative primary antibody control and SRRM2 at 0 dpi and 7 dpi (middle), with intronic versus exonic signal highlighted (bottom). (C) Heatmap of intron-level differentially enriched genes (DEGs) from ARTR-seq, grouped by nuclear speckle enrichment at 0 dpi, shared across 0 and 7 dpi, or enriched at 7 dpi, with selected genes labeled. (D) Representative 3D Airyscan images of MuSC nuclei at 0 dpi and 7 dpi stained for Myod1 RNA puncta (magenta) and SRRM2-defined NS (green). Scale bar, 5 μm. (E) Violin plot of the minimum distance from each *Myod1* RNA punctum to the nearest nuclear speckle for 0 dpi and 7 dpi (all RNA puncta from three animals per group, 18–28 cells per animal, pooled by condition). (F) Density plots of junction-weighted splicing percentage for genes with or without significant nuclear speckle intron enrichment (NS+ and NS−), stratified by gene expression bin (low, mid, high) and condition (0 dpi, 7 dpi). FDR values for NS+ versus NS− comparisons are indicated. (G) Genome browser view of *Ccnd1* showing long-read–defined isoforms, ARTR-seq enrichment, and domain architecture across 0 dpi and 7 dpi, with the dashed box marking a novel 3′ splice site detected in both conditions. (H) Bar plots of *Ccnd1* total gene expression, isoform-specific expression, and isoform fractions for isoforms 1 and 2 at 0 dpi and 7 dpi. (I) Bar plot of mean AlphaFold2 pLDDT for Ccnd1 isoforms 1 and 2. (J) Predicted 3D protein structures for CCND1 isoforms 1 and 2 generated from AlphaFold2.

Gene-level and exon-level ARTR-seq analyses showed related but not identical patterns of nuclear speckle enrichment across regeneration (**Fig. S3D**). At the gene level, enriched targets were broadly distributed across 0 dpi, shared, and 7 dpi categories, with quiescence-associated genes such as *Notch1* and *Pcna* enriched at 0 dpi, and activation-associated genes such as *Myh6*, *Tbx1*, and *Notch3* enriched at 7 dpi. In contrast, exon-level enrichment was more strongly skewed toward 7 dpi, consistent with increased exon-associated nuclear speckle interactions during activation. Notably, *Malat1* was shared across both states in the gene-level dataset and remained shared at the exon level, as expected for a canonical speckle-associated RNA. We also detected *Meg3* among the shared enriched transcripts in both datasets, consistent with prior work showing that MEG3 lncRNA can associate with NS^42^. Together, these analyses indicate that while intron-level enrichment most strongly captures state-specific nuclear speckle-associated transcript programs, gene- and exon-level enrichments also preserve known nuclear speckle RNAs and reveal a broader shift toward exon-associated interactions during MuSC activation. Gene-ontology analysis across gene-, intron-, and exon-level datasets each suggested genes associated with RNA splicing, RNA metabolic processes, RNA-processing, DNA replication, cell-cycle regulation and chromatin-linked functions (**Fig. S3E**). These results are consistent with previous ARTR-seq^19^ and APEX-Seq^43^ datasets and suggest NS interact with transcripts involved with quiescence, myogenic differentiation, splicing machinery and progression through the cell cycle after injury.

We next examined how nuclear speckle association relates to splicing completion as a function of expression level. Using the junction-weighted splicing percentage, we compared nuclear speckle-enriched intron targets (NS+) with non-nuclear speckle genes (NS−) after binning genes into low, mid, and high expression groups (**Fig. 3F**). For transcripts with low expression, NS− genes showed slightly higher splicing percentages than NS+ genes at both 0 dpi and 7 dpi. Conversely, for genes with high expression, NS+ genes were shifted toward nearly fully spliced transcripts at both time points, with a stronger separation at 7 dpi. These data indicate that nuclear speckle-associated intronic transcripts that are robustly expressed are preferentially linked to enhanced splicing completion, which is in line with previous TSA-Seq measurements^9^. For example, we detected cyclin D1 (*Ccnd1*) ^44^ enriched in NS in both 0 and 7 dpi MuSCs, with ARTR-seq enrichment mapping to regions that overlap intron-containing long-read isoforms across both conditions (**Fig. 3G**). Upon injury, total *Ccnd1* gene expression increased markedly (**Fig. 3H, left**), accompanied by a significant shift in isoform usage toward isoform 1, which contains an extended 3′UTR (∼2.7 kb) relative to the quiescent-dominant isoform 2 (∼445 nt 3′UTR) (**Fig. 3H, right**). In addition, isoform 2 uses a novel 3′ splice site in the terminal region, generating a truncated transcript that removes the downstream coding sequence in isoform 1. By contrast, isoform 1 produces a slightly shorter protein (lacking 22 C-terminal amino acids), retains both Cyclin-C and Cyclin-N functional domains and displays comparable predicted structural confidence (pLDDT) (**Fig. 3I, J**), suggesting that the switch does not disrupt core protein folding. Rather, the extension of the 3′UTR in the injury-dominant isoform provides an expanded platform for post-transcriptional regulation, potentially enabling finer control of *Ccnd1* expression and function during the regenerative response. More broadly, the overlap between transcripts identified by ARTR-seq and RNA-seq splicing changes suggests that a subset of regeneration-associated splicing events occur in RNAs that are also proximal to NS, consistent with a regulatory relationship at the global level.

### Old aged muscle stem cells display reductions in nuclear speckle size and numbers and increased splicing compared to young muscle stem cells

Old aged MuSCs are known to have activation defects and impaired tissue regeneration^45,46^, but there is no information for how NS are altered in old age. We first assessed if NS changed in old age (**Fig. 4A**), and isolated MuSCs from hind limb muscles of old aged (24-27 months) mice. We immuno-stained for SRRM2 and Lamin A/C and performed 3D super-resolution microscopy, as above (**Figs. 4B-D**). We observed ∼28% decreases in the number and ∼8% decrease in the diameter of NS when compared to young MuSCs (**Figs. 4C-D**). These results show old aged MuSCs lose both nuclear speckle numbers and size.

**Figure 4.**
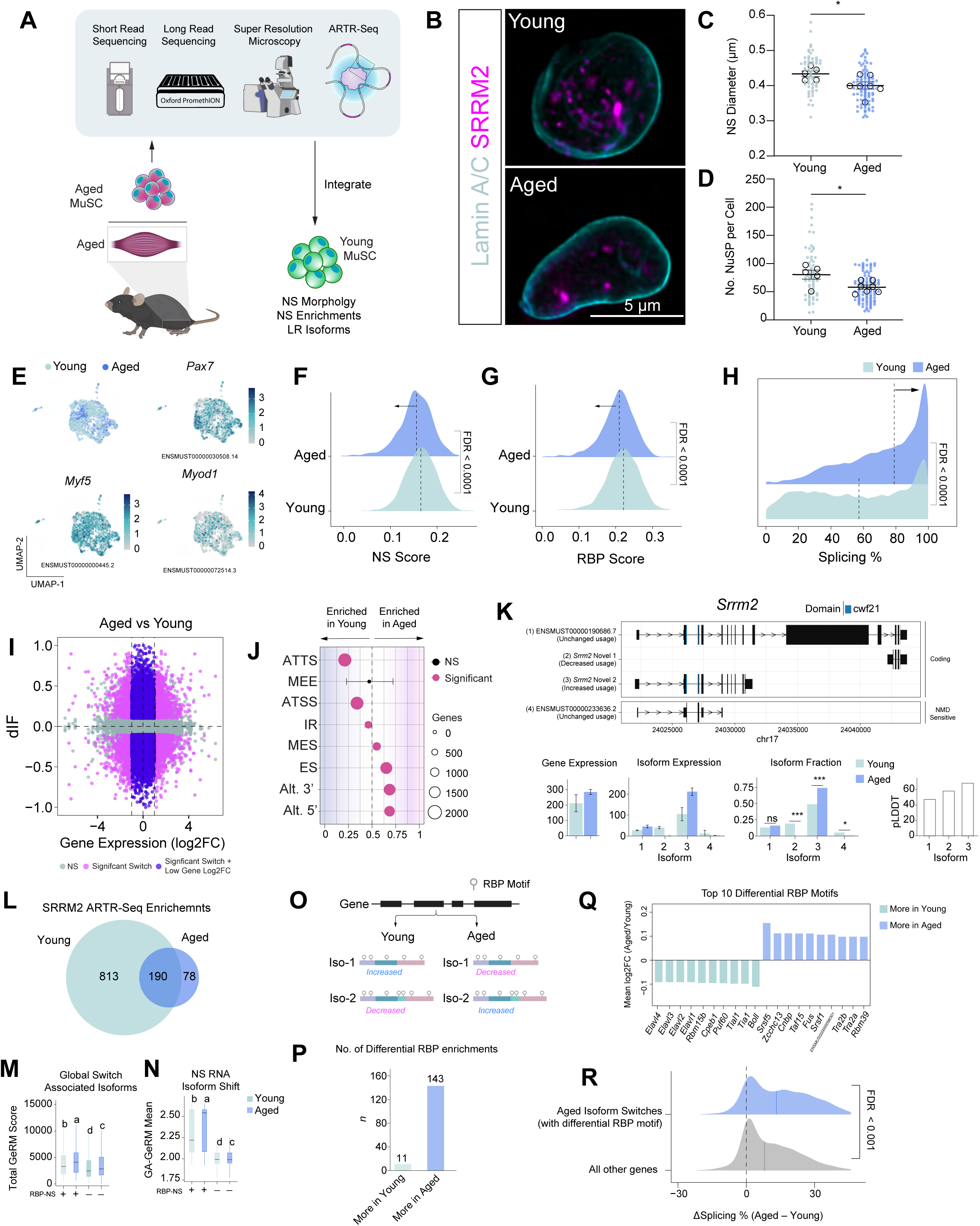
Aging alters nuclear speckle organization, nuclear speckle–RNA binding protein expression, and splicing programs in MuSCs. (A) Schematic of experimental workflow integrating short-read and long-read single-cell RNA-seq, super-resolution imaging of NS, and ARTR-seq in young and aged MuSCs. (B) Representative Airyscan images of NS in young (top) and aged (bottom) MuSC nuclei stained for Lamin A/C (cyan) and SRRM2 (magenta). Scale bar, 5 μm. (C) Quantification of the volume of individual NS in young and aged MuSCs (average nuclear speckle volume per cell). Each point represents a single cell (10–17 cells per biological replicate), with 5–7 animals per group; bars indicate mean ± SEM. (D) Quantification of the number of NS per nucleus in young and aged MuSCs (average nuclear speckle number per cell). Each point represents a single cell (10–17 cells per biological replicate), with 5–7 animals per group; bars indicate mean ± SEM. (E) UMAP of integrated MuSCs from young and aged muscles colored by age group, with feature plots showing long-read isoform expression of *Pax7*, *Myf5*, and *Myod1*. (F,G) Density plots of nuclear speckle (NS) score and RNA binding protein (RBP) score in Young and Aged MuSCs, with median values indicated by dashed lines and FDR values shown. (I) Density plot of junction-weighted splicing percentage in young and aged MuSCs, with median values indicated by dashed lines and FDR shown. (J) Scatterplot of Δ isoform fraction (dIF) versus gene-level log₂ fold change (aged versus young) for all tested isoforms, with points colored by switch significance and gene-level change category. (K) Dot plot summarizing significant isoform switches grouped by alternative splicing class in young or aged MuSCs; x axis indicates enrichment in Young or Aged, point size reflects the number of genes, and color denotes significance. (L) Genome browser view of *Srrm2* showing long-read–defined isoforms, annotated domains, and bar plots of total gene expression, isoform-specific expression, isoform fractions, and mean AlphaFold2 pLDDT values for representative *Srrm2* isoforms in young and aged MuSCs. (M) Venn diagram showing overlap of SRRM2 ARTR-seq enriched transcripts in Young and Aged MuSCs. (N) Boxplots of total generalized RNA multivalency (GeRM) scores for globally switched isoforms stratified by RBP–NS gene membership and age group. (O) Boxplots of mean GA-rich GeRM scores for isoform switches within the NS RNA gene set in Young and Aged MuSCs. (P) Schematic of differential isoform usage within a gene and corresponding changes in RNA binding protein motif content between Young and Aged states. (Q) Bar plot showing the number of differential RNA binding protein motif enrichments that are greater in young or greater in aged isoform switches. (R) Bar plot of the top differential RNA binding protein motifs enriched in young or aged isoform switches, shown as mean log₂ fold change (Aged/Young). (S) Density plot of ΔSplicing% (Aged − Young) for isoform switches containing differential RNA binding protein motifs compared with all other genes, with median values indicated and FDR shown.

To determine whether old aged MuSCs exhibit reduced expression of *Srrm2* and other nuclear speckle-associated RNA binding proteins, we performed paired short-read and long-read scRNA-seq on mononuclear cells isolated from biological replicates of uninjured hindlimb muscle from old aged mice (n = 2; 27-months), as above (**Figs. S4A and S4B**). We then re-clustered MuSCs from young and aged muscles (**Fig. 4E**) and confirmed the expected distribution of canonical myogenic isoforms across the MuSC manifold (**Fig. 4E**). Since age-associated declines in RNA binding protein and splicing factor expression have been reported across multiple tissues^47,48^, we quantified genes annotated to nuclear speckle and RNA binding protein Gene Ontology categories. We found that both genes in NS and RNA binding proteins were modestly but significantly reduced in old aged MuSCs relative to young MuSCs (**Figs. 4F-G**). Differential expression analysis identified 82 downregulated genes including several Rbm family splicing regulators (*Rbm4b*, *Rbm10*, *Rbm25*, *Rbm27*, and *Rbm39*) and 59 upregulated genes including *Srrm2*, *Srrm1*, and *Snrpb2* in old aged MuSCs (**Fig. S4C**). Together, these data suggest that aging remodels, rather than depletes the nuclear speckle and RNA binding protein network.

The attenuation of the number and size of NS in old aged MuSCs along with small differences in expression suggest changes in splicing. Using the junction-weighted splicing percentage defined above, we compared global splicing distributions between young and old aged MuSCs (**Fig. 4H**). Old aged MuSCs showed broadly elevated splicing percentages, with a distribution that more closely resembles activated 7 dpi MuSCs than uninjured young cells. This change is consistent with prior cross-species analyses^47^ showing that aging is accompanied by an increased ratio of spliced exon junction reads to unspliced junction reads, reflecting fewer unspliced transcripts with age. To focus on regulation of shared genes, we calculated ΔSplicing% and found many shared transcripts are more completely spliced in old aged MuSCs (**Fig. S4D**). To further confirm changes in splicing, we assessed changes in isoform usage independent of gene-level abundance and identified 5,331 significant isoform switches across 3,245 genes (FDR < 0.05, |dIF| ≥ 0.1) independent of changes in gene expression (**Fig. 4I**). Annotation of these switches by alternative splicing class showed that old aged MuSCs were enriched for isoforms containing mutually exclusive exons, skipped exons, and alternative 3′ and 5′ splice sites (**Fig. 4J**). Notably, the nuclear speckle scaffold gene Srrm2 underwent an isoform switch in old aged MuSCs despite a non-significant change in overall gene expression (**Fig. 4K**). In old aged MuSCs, isoform usage shifted toward an isoform lacking the large exon that encodes the intrinsically disordered region of SRRM2 and was predicted to have a higher pLDDT score (**Fig. 4K**). This suggests that the isoform with greater predicted condensate-forming potential is used less frequently in old aged MuSCs. Together, these data indicate that old aged MuSCs shift toward highly spliced, but potentially less tightly regulated, transcript processing programs that are accompanied by changes in protein structure and function.

### Interactions between RNAs and nuclear speckles are reduced in old aged muscle stem cells and associate with alterations in RNA multivalency

A recent study^49^ showed that phase separation behavior of condensation-prone proteins could be modified via interactions with their own multivalent mRNAs. To test if age-associated isoform switching in MuSCs could reshape RNA– nuclear speckle interactions by altering RNA multivalency, we performed SRRM2 ARTR-seq^40^ from old aged MuSCs (n=24-26 months) and called enrichments over the negative primary antibody background, as above (**Fig. S4E**). Old aged MuSCs exhibited a marked reduction in RNAs associated with NS, retaining 268 of 1,003 (27%) of the enriched genes in young MuSCs (**Fig. 4L; Fig. S4F**). Young MuSC NS were enriched for transcripts encoding nuclear speckle and splicing proteins (*Srrm1, Srsf6, Srsf7, Srsf10*), as well as quiescence-associated genes (*Notch1*, *Ccnd1*), which were absent in old aged MuSC ARTR-seq libraries (**Fig. S4F**). We next asked whether isoform switching with age altered the multivalent potential of RNAs. We quantified total generalized RNA multivalency (GeRM) scores for each isoform switch, weighted by isoform expression in each condition. We found that multivalency was highest in isoforms encoding RNA binding proteins and NS, including *Srrm2* (Fig. S4G), and increased in old aged MuSCs compared to young (**Figs. 4M-N**). Notably, this pattern was not observed after injury (**Fig. S4H**), indicating that enhanced RNA multivalency is a feature of aging rather than a consequence of activation. These results indicate aging results in increased production of RNAs with sequence features that should favor condensate-linked regulation, but the reduced size and abundance of SRRM2-positive speckles may limit how effectively those RNAs are captured and spliced within speckles. Together, these results suggest that old age is associated with reduced RNA colocalization with NS despite increased multivalency of isoforms.

To understand why increases in multivalency in old aged isoform switches occurred, we evaluated transcript architecture. We detected that old aged isoforms displayed greater exon numbers and significantly longer lengths^50^ (a median increase of 394 nucleotides) when compared to young (**Fig. S4I**). The total number of RNA binding protein motifs also scaled strongly with transcript length across switched isoforms (**Fig. S4J**), and the number of differential RNA binding protein motifs were increased in old age compared to young (**Fig. 4P**). Representative motifs enriched in old aged isoforms included *Srsf1*, which suppresses autophagy^51^ that is critical for MuSC activation^52^, as well as TRA2 proteins that ferry multivalent RNAs to NS^49^. These RNA binding proteins are commonly found within and near NS, but their RNAs were not enriched in SRRM2 ARTR-seq datasets. In contrast, motifs enriched in young MuSCs included *Elavl1*, (HuR) which bind to 3’ UTRs and increases mRNA translation^53^ and stability^54^ (**Fig. 4Q**). Isoform switches containing differential RNA binding protein motifs also exhibited significantly higher ΔSplicing% values than the background gene set (**Fig. 4R**). Taken together, these data suggest that although old aged MuSCs show fewer RNA interactions with NS, they undergo isoform switches within the same genes toward longer, more multivalent transcripts with increased RNA binding protein motif content.

## Discussion

Herein, we build an understanding of how NS and RNA processing^55,56^ influence the regenerative function of MuSCs. We map splicing behavior during regeneration and in old age, which demonstrated pathological interruptions to RNA flow observed in youth. Our construction of an isoform-resolved and splicing atlas in regeneration and aging combined with subnuclear maps of RNA interactions with NS and predicted protein folding offer a new resource to interpret stem cell behavior and consequences of splicing alterations.

RNA binding proteins are critical regulators of RNA fate and influence trafficking to different cellular sites^57^, promotion of decay^58^ and translation into proteins^59,60^. Profiling SRRM2, an RNA binding protein in NS, in MuSCs before and after muscle injury revealed increases in the size and number of SRRM2 condensates. Loss of SRRM2 impinged on MuSC activation and transplantation of MuSCs with SRRM2 loss displayed reductions in myofiber size. These results were in line with old aged MuSCs that displayed loss of SRRM2, and were previously shown to display defects in activation but comparable engraftment efficacy^61,62^ to young MuSCs. Nuclear speckle enlargement has previously been shown to be driven by decreases in transcription or splicing from cell stress^10,12^, which contrasts with our results. To investigate the relationship between MuSC activation and the local nuclear speckle splicing microenvironment, we used long-read RNA sequencing and ARTR-seq in MuSCs. We found intron retention was increased in uninjured MuSCs, while splicing increased after injury and isoforms that switched displayed increased propensity to fold with greater confidence (higher pLDDT). These results are consistent with previous observations that showed adult stem cells utilize intron retention to maintain quiescence^4^, and response to injury / activation promotes utilization of alternative transcription termination sites or polyadenylation sites, which has also been observed in hematopoietic stem cells^63^. The observed changes in open reading frames from increased splicing during MuSC activation may afford MuSCs an energetically efficient mechanism to adjust protein assembly, folding, function and stability^64^ without altering expression or protein synthesis rates. Our data showing isoform shifts with increased protein folding potential further support this model and are also consistent with differentiation of embryonic stem cells into specific cell fates^65^. Uniquely, we found NS shape splicing completion, whereby transcripts that were proximal to SRRM2 showed amplified expression and splicing after muscle injury. These transcripts included regulators of quiescence, RNA processing and chromatin packaging before injury and shifted to myogenic activation and differentiation factors after injury. Integrating these results suggest NS shape splicing that is required for MuSC activation.

Aging is a multifactorial pathological process that impinges on multiple levels of adult stem cell function^45^. The loss of NS in old aged MuSCs and increase in splicing and RNA multivalency show a new dimension through which stem cell regenerative potential is compromised. Several mechanisms may contribute to the loss of NS and dysregulated splicing in old aged MuSCs. First, increases in oxidative stress without injury, and over-activation of kinases that impinge on phase separation of NS may drive reductions in cohesion of nuclear speckle scaffold proteins^16,66^. Multiple age-associated diseases that display exacerbated oxidative stress including amyotrophic lateral sclerosis^67^, cancer^68^, Alzheimer’s disease ^69^, and myotonic dystrophy ^70^ also contain dysregulated NS. Additionally, old aged MuSCs have been shown to display increases in oxidative stress and hyper-activation of kinases^66,71^ such as JAK-STAT^72^, p38-Mitogen-Activated Protein Kinase (p38-MAPK)^73^ and mTORC1^74^, which have been shown to phosphorylate nuclear speckle proteins in their intrinsically disordered regions^16^ and disrupt charge distributions as well as phase separation behavior^75^. Another mechanism which may contribute to loss of NS in old age are changes in the subcellular localization of specific multivalent RNAs that encode for condensation prone proteins^49^. Our observations that old aged NS lose interactions with multivalent RNAs and gain exons as well as RNA binding protein motifs suggest interruptions to the negative feedback loops that maintain protein-RNA condensate size and function^49^. While RBP motif gain does not necessarily imply increased RBP occupancy, as most high-affinity RBP motifs in expressed RNAs are not bound in vivo ^76^, our isoform-switching data remain highly relevant because age-associated exon gains and losses may alter not only the number of RBP motifs present in mature mRNAs, but also the local sequence and structural contexts that determine whether these motifs are available for binding. These changes may create or expose regulatory motifs in mature transcripts while also altering splice-site-proximal and intronic regulatory environments in nascent pre-mRNAs. Third, old aged MuSCs lose histone proteins^46,77^, which reduce transcript processing speed and RNA polymerase II elongation rates^78^ that increase in old age. As a result, old aged MuSCs may experience reduced splicing fidelity and increased splicing noise from faster transcriptional elongation. Future experiments in this space will evaluate how splicing kinetics and transcriptional elongation are modified for genes proximal to NS.

Together, our study identifies NS as a new layer of post-transcriptional control in adult stem cells that link condensate organization to RNA processing, isoform choice, and regenerative output. In old age, this relationship becomes disrupted resulting in rewiring of RNA fate, and pathological isoform states that impair stem cell function. More broadly, this work establishes a framework for understanding how RNA–condensate interactions shape stem cell behavior across regeneration and aging, and provides a foundation for targeting nuclear speckle biology to restore regenerative capacity in old age.

## Methods

**Table 1.** Key Resources.

| Reagent or resource | Source | Identifier |
| --- | --- | --- |
| **Chemicals, and Cell Culture Plates** |  |  |
| Barium Chloride | Sigma-Aldrich | 202738-5G |
| Tamoxifen | Sigma-Aldrich | T2859-1G |
| Dispase II | Sigma-Aldrich | D4693-1G |
| Collagenase II | ThermoFisher | 17101015 |
| DMEM (Dulbecco's Modified Eagle Medium) | Fisher Scientific | 11995065 |
| Matrigel | ThermoFisher | CB-40234 |
| Cell-Tak | Corning | 354240 |
| Poly-L-lysine hydrobromide | Millipore Sigma | P4707 |
| Collagen I, rat tail | ThermoFisher | A1048301 |
| Penicillin-Streptomycin | Fisher Scientific | 15-140-122 |
| Heat inactivated fetal bovine serum (FBS) | ThermoFisher | 10082147 |
| Horse serum | Fisher Scientific | 16050122 |
| DMEM (Dulbecco's modified Eagle medium) | Fisher Scientific | 11995065 |
| F-10 Ham | Sigma Aldrich | N6635 |
| BSA (bovine serum albumin) | Invitrogen | 16140071 |
| EDTA | ThermoFisher | 15575020 |
| bFGF (basic fibroblast growth factor) | Gibco | PHG0263 |
| GSK626616 | R&D Systems | 6638 |
| DPBS (Dulbecco's phosphate-buffered saline) | ThermoFisher | 14040141 |
| HBSS (Hank's balanced salt solution) | Gibco | 14175095 |
| 4% paraformaldehyde | ThermoFisher | J19943.K2 |
| DAPI | ThermoFisher | 62248 |
| Triton X-100 | Millipore Sigma | X100 |
| Goat serum | ThermoFisher | 31873 |
| Donkey serum | Sigma Aldrich | S30 |
| Tween-20 | Millipore Sigma | P9416 |
| Glycine | ThermoFisher | 220911000 |
| Co-Detection Antibody Diluent | ACD | 323160 |
| Chamber cover Glass | Cellvis | C8-1.5H-N |
| 48-well plate with glass bottom | MatTek | P48G-1.5-6-F |
| 96-well plate with glass bottom | Cellvis | P96-1.5H-N |
| 12-well plate | ThermoFisher | 12565321 |
| RNAscope Myod1 probe | ACD | 316081 |
| RNAscope positive probe | ACD | 320881 |
| RNAscope negative probe | ACD | 320871 |
| TSA Vivid Fluorophore 520 (1:1500) | ACD | 323271 |
| Vector Laboratories ImmEDGE™<br>Hydrophobic Barrier Pen | Fisher Scientific | NC9545623 |
| Prolong Diamond Antifade Mountant | Invitrogen | P36970 |
| **Antibodies** |  |  |
| Mouse anti-Pax7 (concentrate) | Developmental<br>Studies<br>Hybridoma<br>bank (DHSB) | Concentrate 0.1 ml |
| Rabbit anti-SRRM2 | Novus<br>Biologicals | NBP2-55697 |
| Rabbit anti-SON | Proteintech | 28046-1-AP |
| Lamin A/C Antibody (E-1) Alexa Fluor® 488 | Santa Cruz<br>Biotechnology | sc-376248 AF488 |
| Chicken anti-GFP | ABCAM | ab13970 |
| Rabbit anti-Laminin 1+2 | ABCAM | ab7463 |
| Mouse anti-MyH3 (supernatant) | DHSB | F1.652 |
| Goat anti-Rabbit IgG (H+L) Cross-Adsorbed<br>Secondary Antibody, Cyanine5 | ThermoFisher | A10523 |
| Goat anti-Mouse IgG1 Cross-Adsorbed<br>Secondary Antibody, Alexa Fluor™ 555 | ThermoFisher | A-21127 |
| Goat anti-Rabbit IgG (H+L) Cross-Adsorbed<br>Secondary Antibody, 594 | ThermoFisher | A32740 |
| Donkey anti-Chicken IgG Cross-Adsorbed<br>Secondary Antibody, 488 | ThermoFisher | A78948 |
| **Critical Commercial Assays** |  |  |
| Satellite cell isolation kit | Miltenyi Biotec,<br>Germany | 130-104-268 |
| α-7 integrin cell isolation kit | Miltenyi Biotec,<br>Germany | 130-104-261 |
| Click-iT™ Plus EdU Cell Proliferation kit | ThermoFisher | C10639 |
| RNease Micro Kit | Qiagen | 74004 |
| SuperScript III First-Strand Synthesis System for RT-PCR | Invitrogen | 18080051 |
| QuantiTect SYBR Green RT-PCR Kit | Qiagen | 204143 |
| AllPrep Micro Kit | Qiagen | 80284 |
| AllTaq Master Mix Kit | Qiagen | 203144 |
| RNAscope Multiplex Fluorescent assay v2 | ACD | 323100 |
| **Experimental Models: Organisms/Strains** |  |  |
| C57BL/6J | University of Michigan | Jackson stock 000664 |
| Pax7<sup>CreERT2</sup>-Rosa26<sup>Tdtomato</sup> | University of Michigan | Jackson stock 017763 crossed with stock 007676 |
| Pax7<sup>CreERT2</sup>-Rosa26<sup>LSL-Cas9-eGFP</sup> | University of Michigan | Jackson stock 017763 crossed with stock 024857 |
| **Deposited Data** |  |  |
| Young and Old Aged Muscle Stem Cell Single Cell Droplet Long-Read and Short-Read RNA-seq Datasets | This Manuscript | GSE317037 |
| Young and Old Aged Muscle Stem Cell ARTR-seq SRRM2 Datasets | This Manuscript | GSE317249 |
| **Software** |  |  |
| R | The R foundational for statistical computing |  |
| Fiji | National Institutes of Health |  |
| CellProfiler | Broad Institute |  |
| Cellpose | HHMI Janelia<br>Research<br>Campus |  |
| IMARIS | Oxford<br>Instruments |  |
| Prism v.10 | Dotmatics |  |

### Animal Model

All animal procedures were approved by the University of Michigan Institutional Animal Care and Use Committee and the Animal Care and Use Review Office of the Medical Research and Development Command, US Army and were in accordance with the U.S. National Institute of Health (NIH) guide for the care and use of animals.

### Muscle Stem Cells (MuSC) Isolation

#### Magnetic-Activated Cell Sorting (MACS)

C57BL/6J wild-type mice (3-5 months and 24-27 months old) were sacrificed with CO_2_, hindlimb muscles were harvested, and tissues were processed as previously described^79^. Briefly, muscles were weighed, minced, and digested (20 ml of digest solution: 2.5 U/ml Dispase II, 0.2% Collagenase II in DMEM per mouse) at 37°C for 1 hour. Pipetting of the tissue with an FBS coated pipette was performed at 30 minutes to further break the muscle pieces. Enzymes were then inactivated with stop solution (20% heat inactivated FBS in F10 media) at a 1:1 ratio of digest solution. To remove fat and tendons from the single cell suspension, digested muscle was filtered through a 70 µm cell trainer and centrifuged at 300xg for 10 min at 4°C. MuSCs were then isolated using the Satellite cell Isolation Kit (Miltenyi Biotec) and further enriched with the anti-integrin α-7 microbeads kit (Miltenyi Biotec) per manufacturer’s instructions.

#### Fluorescence-Activated Cell Sorting (FACS) to isolate TdTomato^+^ MuSCs

To capture MuSCs and their progenitors after muscle injury, Pax7^CreERT2^-Rosa26^Tdtomato^ (3-5 months old) mice were injected with tamoxifen for 5 days and allowed to recover for 2 days. After the recovery period, 1.2% Barium Chloride (BaCl_2_) was injected along the tibialis anterior (TA) muscle for a total of 40 µl. The TA from both hindlimbs was harvested at 7 days post injury (dpi). TA muscles from C57BL/6J wild-type mice were used as controls (unstained, live cells) for fluorescence-activated cell sorting (FACS). The muscles were digested as described above. The single cell suspension was then rinsed with FACS buffer (2% heat inactivated FBS, 5 mM EDTA in HBSS), centrifuged and cell pellet resuspended in 1-2 ml of FACS buffer. DAPI (1 µg/ml) was added to the samples and filtered through a 30 µm cell strainer. Samples were loaded into the Sony MA900 sorter and TdTomato^+^ cells were isolated.

### In vitro *Srrm2* knockdown on MuSCs

MuSCs from wild type mice (3-5 months old, n=4) were isolated via MACS, as above, and plated on a Matrigel-coated 48-well plate at 13,000 cells/cm^2^. Twenty-four hours after cell seeding, SRRM2 knockdown was performed with the Lipofectamine RNAiMAX reagent following manufacturer’s instructions. Small interference RNA from IDT (mm.Ri.Srrm2.13.1) at 60 nM was used. Lipofectamine control was included to assess any potential effects of the transfection protocol. MuSCs were incubated with the lipofectamine:siRNA complexes for 48 hours. To confirm the knockdown, MuSCs were trypsinized, centrifuged, and resuspended in RLT buffer. RNA was then isolated following the RNeasy Micro kit. Complementary DNA was synthesized following the SuperScript III First-Strand kit. Quantitative RT-PCR was performed with the QuantiTect SYBR Green PCR kit with *Srrm2* (Mm. PT.58.28432809) and Rplp0 (Mm.PT.58.43894205) primers and the plate was run on a QuantStudio 3 instrument. Comparative ΔΔCT values were then analyzed. Immunocytochemistry was also performed on cultured MuSCs on glass bottom well plates to confirm knockdown of *Srrm2* at the protein level. In addition, to characterize potential morphological changes of other key scaffolding proteins in NS, staining of SON was performed, as described below.

To analyze the effect of *Srrm2* knockdown on MuSC state, we first performed Pax7 and EdU staining to assess quiescence and proliferation, respectively, as described below. To further assess the functionality of MuSCs, cells were cultured on 12-well plate (13,000 cells/cm^2^), and *Srrm2* knockdown was performed as described above. Transfection media was removed after 48 hours and cells were incubated with fresh growth media for 5 hours. Then, MuSCs were incubated with differentiation media (5% Horse serum + 1% Pen-Strep in DMEM) for 3 days to induce fusion. Myotubes were fixed and stained as described below.

### In vitro treatment of GSK626616

MuSCs from wild type mice (3-6 months old, n=3-4) were isolated via MACS and plated at 15,000 cells/cm^2^ on chamber cover slides coated with poly-lysine and collagen. Coating was prepared by incubating the cover slides with 100 µg/ml of poly-l-lysine for 30 minutes at RT, rinsed 3x3 minutes with distilled water, air dried for another 30 minutes, incubated with 50 µg/ml of collagen diluted in 20 mM acetic acid for 1 hour at RT, and rinsed 3x3 minute with PBS. Twenty-four hours after cell seeding, MuSCs were treated with DMSO or 5 µM of GSK626616, a selective DYRK inhibitor, for 2 and 24 hours, to assess changes in nuclear speckle morphology and MuSC activation, respectively.

### In vivo CRISPR-Cas9 Gene Knockout of SRRM2 in MuSCs

To target CRISPR-Cas9 gene editing to MuSCs, a Pax7-Cas9 mouse strain was generated by crossing a Pax7^CreERT2^ driver mouse strain (JAX# 017763) with a Rosa26-loxP-Stop-loxP-Cas9-eGFP reporter mouse strain (JAX# 026175). AAVs targeting SRRM2 were manufactured by VectorBuilder^TM^. The pAAV plasmid backbone was designed with two guide RNAs (CTTCGAGATCGTCGTCGTGC and GATCGAGAGCTCCGGCGTCG) targeting SRRM2, a scaffold sequences specific to S. Pyogenes Cas9 (GTTTTAGAGCTAGAAATAGCAAGTTAAAATAAGGCTAGTCCGTTATCAACTTGAAAAAGTGGCACCGAGTCGGTGC), a U6 promoter, ampicillin resistance gene for bacterial selection of transformed E. coli, and a mCherry reporter under a CAG promoter to assess virus transduction. Another pAAV plasmid encoding a non-targeting/scramble gRNAs (GTGTAGTTCGACCATTCGTG and GTTCAGGATCACGTTACCGC) was used to control for nonspecific effects of CRISPR-Cas9 delivery. Plasmid DNA was isolated from transformed E.coli and packaged into AAV9 viral particles.

To assess the efficiency of transduction, AAVs for SRRM2 KO and scramble control were thawed on ice, resuspended in sterile PBS and 20 µl (3e11 vg) and 30 µl (4.5e11 vg) were injected into TA and gastrocnemius muscles, respectively (n=18-19 mice/group). Three weeks after injection, muscles were harvested and GFP^+^mCherry^+^ MuSCs were isolated via FACS, as previously described^46,80^.

To validate CRISPR-mediated gene editing, DNA was isolated from mCherry^+^ cells using the AllPrep Micro kit following the manufacturer’s instructions. The genomic region flanking the CRISPR target site within SRRM2 was amplified by PCR for both gRNAs (Forward: GGTCTAGGACACGATCACCAGTTC, Reverse: GGCTGAGGAGAATTTGAACGAC, expected size 589 bp and Forward: CCCCACAGGAAAGAAGTGAGT, Reverse: CTGGACCGCCGATGAGTAG, expected size 976 bp) using the AllTaq Master Mix Kit. PCR products were submitted for purification and further Sanger Sequencing for forward and reverse primers (Eurofins Genomics). Chromatograms were analyzed using Benchling.

For transplantation experiments, AAVs were injected into TA and gastrocnemius muscles as described above (n=6-7 mice/group) and GFP^+^mCherry^+^ MuSCs were isolated 3 weeks after injection. The day before isolation, TA muscle from both hindlimbs of wildtype mice (n=6-7) were injected with BaCl_2_. Isolated MuSCs from SRRM2 KO and scramble groups, were injected into the TA of each hindlimb, respectively. Tissues were harvested 21 days post-injury, embedded in Optimal Cutting Temperature (OCT) and flash-frozen in isopentane pre-chilled in liquid nitrogen. Muscle cross-sections were obtained along the length of the muscle (proximal, middle, and distal). Sections were then fixed in cold acetone at -20°C for 10 minutes, air-dried for 10 minutes, outlined with hydrophobic pen, and rinsed with 1XPBS. Samples were then incubated with blocking buffer (10% donkey serum in PBS) for 1 hour, followed by overnight incubation at 4°C with a rabbit anti-laminin 1+2 antibody (1:500) and a chicken anti-GFP antibody (1:1000) diluted in 5% donkey serum in PBS. Cross-sections were then rinsed with 1X PBS 3X for 5 minutes, incubated for 2 hours at RT with goat anti-rabbit Alexa Fluor 594 (1:500) and donkey anti-chicken Alexa Fluor 488 (1:500). Samples were rinsed with 1XPBS 3X for 5 minutes, incubated with DAPI (1:1000) for 10 minutes, and rinsed again with 1XPBS. Cover slides were mounted with prolong diamond antifade mountant.

Images were acquired with a Zeiss Laser Scanning Microscope (LSM) 900 Airyscan at 20X with confocal mode. Outlines for fibers were obtained using a customized model in Cellpose with the cyto3 model as a base. A non-injected muscle was used as a negative control to determine the threshold of GFP^-^ fibers. In addition, the mean fluorescence intensity (MFI) of ∼10 GFP^-^ fibers was measured and the average + 2* standard deviation was calculated to obtain the threshold value of GFP^+^ fibers per cross-section. The GFP MFI of individual fibers was obtained in Fiji.

### RNA-SCOPE

Uninjured and BaCl_2_-injured muscles (7dpi) were harvested from wild type mice (3-6 months, n=3/group) and MuSCs were isolated via MACS. Cells were seeded on a poly-l-lysine and collagen-coated 96-well plate as described above and allowed to attach overnight. MuSCs were then processed with the RNAscope ® Multiplex Fluorescent Kit v2 following the manufacturer’s instructions for RNAscope assay combined with Integrated Co-Detection Workflow. Briefly, cells were fixed with 4% PFA for 30 minutes at RT, rinsed 3X with 1X PBS, and dehydrated and rehydrated with 50-100% ethanol solutions for 1 minute. Samples were then incubated with Hydrogen Peroxide for 10 minutes at RT, rinsed 3X with distilled water, and incubated overnight at 4°C with SRRM2 antibody (1:200) diluted in co-detection antibody diluent. Cells were rinsed 3X with 0.1% tween-20 in PBS (PBS-T), fixed again in 4% PFA for 30 minutes at RT, and rinsed 3X with PBS-T. Samples were then incubated with protease III (1:15) for 10 minutes at RT on a humidity control tray, rinsed 3X with 1X PBS. Target mRNA was hybridized with *Myod1* probe for 2 hrs at 40°C in the HybEZ^TM^ Oven and rinsed 3X with wash buffer. Probe was then sequentially hybridized with amplification molecules to increase RNA signal, followed by sequential incubation of TSA® Vivid-labeled probe to bind to the amplification trees through a series of horseradish peroxidase reactions. To visualize SRRM2 signal, MuSCs were incubated with Alexa Fluor 647 goat anti-rabbit diluted in co-detection antibody solution (1:200) for 30 minutes at RT. Samples were rinsed with PBST-2 3X for 2 minutes, incubated with DAPI for 5 minutes, and rinsed 3X with PBS. Positive and negative control probes, which include housekeeping genes (POLR2A and PPIB) and bacterial gene (dapB), respectively, were used to validate the reliability of the assay. In addition, RNase treatment was also performed to assess the specificity of the RNA signal.

Z-stacks were acquired with a Zeiss LSM 900 Airyscan2 with the super-resolution mode at 63X with 3X digital magnification. Images were analyzed with IMARIS software. Surface segmentation was performed for DAPI, SRRM2, and RNA puncta signal. DAPI segmentation was used to isolate RNA puncta signal in the nucleus and the shortest distance of RNA puncta to SRRM2 signal was then obtained with IMARIS.

### Immunostaining and Imaging Analysis

To characterize the morphology of NS in uninjured and injured MuSCs, the cells were seeded on chamber cover glass pre-coated with Cell-Tak and fixed (4% PFA for 20 minutes) immediately after isolation. Cells were then permeabilized with 0.5% triton X-100 for 10 minutes at room temperature (RT), washed twice with 0.1% triton X-100 in PBS and blocked with 4% BSA in PBS for 30 minutes at RT. Overnight 4°C incubation of SRRM2 primary antibody (1:200 diluted in 3% BSA in PBS) was followed by three rinses with 0.1% triton X-100 in PBS and incubation of secondary antibody (Goat anti-Rabbit Cyanine 5, 1:200 diluted in 3% BSA in PBS) for 2 hours at RT. Samples were also incubated with Lam A/C conjugated to Alexa Fluor 488 (1:50) to visualize nuclear envelope. Samples were then rinsed three times with PBS, incubated with DAPI (1:1000) for 10 minutes and rinsed again with PBS. Single-cell images were then obtained with a Zeiss LSM 980 Airyscan 2 with the super-resolution mode at 60X with 5X digital magnification. Airyscan processing was performed with the Zeiss software. Three-dimensional measurements of NS were analyzed with a custom macro in Fiji.

To assess proliferation, MuSCs were incubated with EdU (10 µM) a day before fixation. Then, EdU was labeled through a click-chemistry reaction following manufacturer instructions. MuSCs were then rinsed with PBS, permeabilized with 0.2% triton X-100 in PBS for 15 minutes, rinse again with PBS, and block (2% BSA, 5% goat serum in 0.1% tween-20) for 1 hour. Primary antibodies (Pax7- 1:100, SRRM2-1:200) were diluted in 1% BSA in 0.1% tween-20, added to the samples and incubated overnight at 4°C. Samples were then rinsed with 0.1% tween-20 and incubated with secondary antibodies (Pax7-Alexa Fluor goat anti-mouse 555 and SRRM2 Alexa Fluor goat anti-rabbit 647) for 2 hours at RT. Samples were then rinsed with PBS and incubated with DAPI (1:1000) for 10 minutes and rinsed again with PBS. For Pax7 and EdU, images were obtained at 20X across the wells. The corrected mean fluorescence intensity (Raw intensity- randomly selected regions of interest in background) for Pax7 was obtained across the field of views. EdU was quantified as the percentage of EdU^+^ MuSCs divided by the total MuSCs. Single cell images were obtained with a CSU-W1 SoRa (Yokogawa Spinning Disk Field Scanning Confocal System) at 100X with 2.8X digital magnification. Deconvolution was performed using the Richardson-Lucy method with the Nikon software. The mean fluorescence intensity of SRRM2 was obtained per nuclei with previous background subtraction using the rolling ball radius method in Fiji.

To characterize another NS scaffolding protein (SON) with and without SRRM2 knockdown, MuSCs were fixed as described above, permeabilized with 0.5% triton X-100 in PBS for 10 minutes at RT, rinsed twice with PBST and blocked for 1 hour at RT (2% BSA + 5% goat serum in PBST). Samples were then incubated with SON antibody (1:1000 diluted in 1% BSA in PBST) overnight at 4°C, rinsed 3X with PBST and incubated with secondary antibody for 2 hours at RT (Alexa Fluor goat anti-rabbit 647, 1:1000 diluted in 1% BSA in PBST). MuSCs were rinsed 3X in PBS, incubated with DAPI (1:1000) for 10 minutes and rinsed again with PBS. Single-cell images were acquired with LSM 900 Airyscan at 63X objective with super-resolution mode. Airyscan processing was performed with the Zeiss software. Three-dimensional measurements of SON signal were then obtained with a customized pipeline in CellProfiler and Fiji.

To evaluate the functionality of MuSCs after SRRM2 knockdown, fixed myotubes were blocked for 1 hour (1% BSA, 22.52 mg ml^-1^ glycine in PBS) and incubated overnight at 4°C with Myh3 antibody (1:20 in 1% BSA in PBST). Then, samples were rinsed 3X with 1XPBST and incubated with secondary antibody for 2 hours at RT (Alexa Fluor-488 goat anti-mouse IgG1- 1:50). Myotubes were imaged with the LSM 900 Airyscan at 10X with camera mode. The fusion index was manually quantified as the percentage of nuclei inside myotubes, with at least two nuclei, divided by the total number of nuclei per field of view.

### Paired Single Cell Short Read and Long Read Sequencing

Single-cell suspensions were obtained from young uninjured (n = 2), young 7 days post-injury (7dpi; n = 2), and aged (n = 3) skeletal muscle using fluorescence-activated cell sorting (FACS) for live cells (PI-negative). Library preparation was performed using the 10x Genomics Chromium Single Cell 3’ Gene Expression HT v3.1 chemistry, targeting 20,000 cells per library. Following cDNA amplification, a portion of each library was reserved for full-length cDNA sequencing using Oxford Nanopore Technologies (ONT). Full-length cDNA was prepared using the ONT Ligation Sequencing Kit V14 (SQK-LSK114) and PCR expansion module (EXP-PCA001), then sequenced on the PromethION platform. A separate portion of cDNA was processed using standard 10X Genomics protocols for short-read sequencing and sequenced on an Illumina NovaSeq X system, targeting a depth of 25,000 reads per cell.

### Single Cell Data Processing

Short-read FASTQ files were aligned to the GRCm39 reference genome using Cell Ranger v8.0.0 (10x Genomics). To minimize ambient RNA contamination, DecontX was applied to the raw count matrices. Cells expressing fewer than 200 genes or with >10% mitochondrial RNA content were excluded. Putative doublets were identified and removed using DoubletFinder ^81^. Downstream processing—including normalization, scaling, identification of variable features, dimensionality reduction, and clustering—was performed using Seurat^82^ (v5) with default parameters unless otherwise noted. Batch correction across samples was carried out using Harmony ^83^, and clustering was performed using the Louvain algorithm at a resolution of 0.2. Cell type identities were assigned based on canonical marker gene expression.

ONT raw pod5 files were basecalled using Dorado v7.3.11. Reads with a minimum Q-score of 9 and length ≥200 bp were retained. Resulting FASTQ files were demultiplexed using the find_barcode() function in FLAMES^84^, which assigns each read to a cell barcode and UMI derived from the matched short-read barcode list. Reads were deduplicated by collapsing on shared barcode–UMI combinations, retaining only the longest representative read from each duplicate group. Long read alignment to the GRCm39 reference genome was performed using minimap2^85^ (v2.26), and isoform detection and quantification were carried out using IsoQuant^34^. Long-read count matrices for both isoform- and gene-level expression were generated and grouped by cell barcode. These matrices were subsequently integrated into the Seurat object as independent assays, enabling joint analysis with matched short-read expression data.

### Splicing percentage from nanopore long-read assignments

Per-read assignments were generated with IsoQuant^34^ on Oxford Nanopore cDNA reads, which outputs a tabular file per sample containing gene and transcript identifiers, a high-level read classification tag, and event tokens indicating features such as intron retention, extra introns, full splice matches, and incomplete splice matches. A custom Python workflow streamed each file, parsed the classification and event annotations, mapped Ensembl gene identifiers to symbols using a GRCm39 GTF, and collapsed read-level information to gene-level summaries within each sample.

To avoid inflating completeness in the presence of intronic signal, any read with intron evidence was treated as unspliced. Specifically, reads were assigned to the unspliced bin if the assignment type indicated genic intron or if event tokens contained intron_retention or extra_intron. All other reads were candidates to contribute fractional evidence of splicing For each non-intronic read we computed a per-read spliced weight in [0, 1]. Full-splice-match reads received weight 1. Mono-exon matches received weight 1 only if the gene is truly single-exon in the reference annotation, defined as all annotated transcripts having exactly one exon; otherwise their weight was 0. For incomplete-splice-match and novel junction classes we quantified partial evidence directly from the exon-block string emitted by the assigner: the observed junction count was the number of exon blocks minus one. The expected junction count was taken from the matched transcript when an isoform identifier was present; otherwise it was the maximum junction count among all annotated transcripts of that gene. To avoid over-penalizing large genes that naturally have many junctions, we applied a fixed cap to the expected junction count with C = 10, so the effective expected count was min(expected, 10). The per-read weight was then defined as observed_junctions divided by that capped expected count, clipped at 1. For example, if a transcript has 20 junctions and a read confirms 2 junctions, the cap sets the effective expected count to 10, giving a weight of 0.2 rather than 0.1; a read confirming 5 junctions gives a weight of 0.5. This cap keeps the metric comparable across genes with widely varying exon counts, noting that many mouse genes have fewer than a dozen exons.

Within each sample and gene we summed the per-read spliced weights to obtain a weighted spliced count and we summed the hard unspliced reads to obtain the unspliced count. We reported the weighted splicing percentage as 100 × weighted_spliced / (weighted_spliced + unspliced). Reads that were neither hard unspliced nor assigned a positive spliced weight were tracked as ambiguous for quality control but were not included in the denominator. This definition treats intron-bearing reads as definitive evidence of incomplete processing, while allowing truncated long reads with informative junction structure to contribute proportional evidence toward completeness. By combining a hard intron guard with a junction-ratio weight and a fixed cap of 10 on the expected junction count, the metric remains robust to systematic truncation and comparable across genes with very different exon counts. A comparative genomics analysis detailing exon–intron organization across mice and humans^86^, substantiated average number of exons per gene is 8.59 for human and 7.58 for mouse, which motivated our cap at 10 used here^87^.

This strategy differs fundamentally from short-read spliced–unspliced frameworks, such as RNA velocity^88^ using velocyto^89^ or kallisto^90^|bustools, which infer splicing status from exon- and intron-compatible fragments in short reads and therefore cannot resolve reads that lie entirely within exons or across long junctions without intronic overlap. Long reads carry explicit exon block structure and isoform identifiers, which enables weighting by the fraction of expected junctions actually observed and avoids reliance on intronic coverage as the sole marker of immaturity. By combining a hard intron guard with ratio-based weights and a cap on expected junctions, the metric yields a gene-level splicing percentage that is robust to systematic truncation, comparable across genes with very different exon counts, and directly grounded in the long-read alignment structure.

### Isoform Switch Analysis

FASTQ pseudobulks for muscle stem cells were generated by subsetting the main single-cell FASTQ files using cell barcodes corresponding to muscle stem cells for each sample. These pseudobulk FASTQs were processed with IsoQuant to generate sample-specific transcriptome GTFs, which were subsequently merged using StringTie ^91^. From the merged transcriptome annotation, a custom FASTA was built with GFFread on the GRCm39 reference genome. Using this custom GTF and FASTA, we constructed a kallisto index in long-read mode (lr-kallisto; kallisto v0.51), which uses an extended k-mer length (k = 63) optimized for long-read pseudoalignment and transcript quantification^92^. We then pseudoaligned the pseudobulk FASTQs to quantify transcript abundance.

The resulting count matrices were imported into R and analyzed using the IsoformSwitchAnalyzeR (v2) workflow^36^. This pipeline incorporates established statistical frameworks for normalization and differential expression at the isoform level^93,94^ and performs isoform switch testing through exon usage and transcript-level abundance estimates^37^. IsoformSwitchAnalyzeR additionally integrates open reading frame prediction and assessment of nonsense-mediated decay (NMD) sensitivity^36,95^ as well as protein domain annotation through the Pfam database^96^. All analyses were performed using default parameters in the IsoformSwitchAnalyzeR pipeline, enabling systematic detection of isoform switches, differential transcript usage, and downstream functional consequences on coding potential and protein features.

### AfphaFold2 structure prediction

To reduce computational time, we employed the small Big Fantastic Database (BFD) option for multiple sequence alignment generation, which uses a reduced subset of the full BFD database. For each protein, the predicted structure with the highest predicted Local Distance Difference Test (pLDDT) score was selected for downstream structural analyses.

### Generalized RNA multivalency (GeRM) analysis

Isoform nucleotide sequences were extracted in FASTA format from the IsoformSwitchAnalyzeR object and included multi-isoform genes identified across the uninjured MuSCs (0 dpi) versus Injury (7 dpi) and Young versus Aged comparisons. Generalized RNA multivalency was quantified using the GeRM framework which assigns a GeRM score to each nucleotide position based on local k-mer sequence similarity. GeRM was run at 5-mer resolution (k = 5) with a multivalency window of 123 and smoothing window of 123. The resulting position-resolved and smoothed k-mer multivalency scores were then processed in R to generate isoform-level summaries. Smoothed GeRM values were aggregated across each transcript to calculate overall multivalency as well as GA-enriched multivalency metrics. GA-rich 5-mers were defined as sequences containing at least 80% purines (A/G) and were further subdivided into A-rich and G-rich subclasses based on nucleotide composition. Transcript-level features, including total GeRM signal, GA-associated signal, and A-rich GA multivalency, were subsequently integrated with isoform usage estimates from IsoformSwitchAnalyzeR for downstream comparative analyses.

### RBP Motif Scanning and Enrichment Analysis

To investigate RNA-binding protein (RBP) regulatory potential associated with isoform switching, we performed motif scanning across all isoforms detected in each pairwise comparison (Young vs. Aged; 0 dpi vs. 7 dpi). Position weight matrices (PWMs) for 208 mouse RBPs were obtained from the CisBP-RNA database^97^ and converted to RNA alphabet for compatibility with transcript sequences. Full-length nucleotide sequences for all isoforms were extracted directly from the IsoformSwitchAnalyzeR nucleotide FASTA, converted from DNA to RNA, and scanned against all RBP PWMs using the universalmotif package^98^ scan_sequences() function with a *p*-value threshold of 1×10⁻⁴ on the forward strand only, yielding per-isoform, per-RBP motif hit counts. A per-isoform valency score was defined as the total number of PWM motif hits across all RBPs, deliberately without length normalization, as increased transcript length in aged isoforms represents a biological signal of interest rather than a technical confound. For gene-level analysis, isoform motif scores were aggregated as expression-weighted means using isoform-level expression values ensuring that highly expressed isoforms contributed proportionally to the gene-level score.

Differential RBP motif enrichment between conditions was assessed for all genes and restricted to significantly switched genes (isoform switch FDR < 0.05, |ΔIF| ≥ 0.1) using paired t-tests across per-gene weighted mean motif scores, with Benjamini-Hochberg correction for multiple comparisons across all RBPs tested. Fold-change was computed as log2((mean condition 2 score + 1) / (mean condition 1 score + 1)) from group means, with a pseudocount of 1 appropriate for count-based motif data. Differential enrichment was assessed using expression-weighted isoform scores, where each isoform’s motif count was weighted by its condition-specific TPM value, ensuring that the gene-level score reflects the RBP binding potential of the isoforms expressed in each condition.

### Reverse transcription–based RNA binding protein binding sites sequencing

ARTR-seq was performed as previously described^40^. Briefly, freshly isolated MuSCs obtained using the MACS protocol (see above) from young uninjured (0 dpi), young 7 dpi, and aged mice were plated onto eight-well chamber slides coated with 10% Matrigel and allowed to adhere overnight. MuSCs were fixed with 1.5% paraformaldehyde (PFA) for 10 min at room temperature, quenched with 125 mM glycine, and permeabilized with 0.5% Triton X-100 on ice for 10 min. Samples were blocked with UltraPure bovine serum albumin (BSA) (1 mg/ml, Thermo Fisher Scientific), stained with SRRM2 antibody at room temperature for 1 hour, and then stained with fluorophore-labeled secondary antibody at room temperature for 30 min. Samples were then incubated with pAG-RTase for an additional 30 min. A reverse transcription reaction mixture was prepared by mixing 2 μM adapter-RT primer (5′-AGACGTGTGCTCTTCC-GATCT-10 N-3′), 0.05 mM biotin-16-dUTP (Jena Bioscience), 0.05 mM biotin-16-dCTP (Jena Bioscience), 0.05 mM deoxythymidine triphosphate (Thermo Fisher Scientific), 0.05 mM dCTP (Thermo Fisher Scientific), 0.1 mM deoxyadenosine triphosphate (Thermo Fisher Scientific), 0.1 mM deoxyguanosine triphosphate (Thermo Fisher Scientific), and RNaseOUT (1 U/μl, Thermo Fisher Scientific) in 50 μl of DPBS supplemented with 3 mM MgCl2. In situ reverse transcription was performed by adding RT reaction mixture to cells and incubating at 37°C for 30 min and then quenched by adding 20 mM EDTA and 10 mM EGTA. To check the success of in situ reverse transcription, cells were stained with streptavidin conjugated with Alexa Fluor 657 (S32357, Thermo Fisher Scientific) and imaged by a Sony LSM900 Airyscan2. The fluorescence intensity distribution on a line was quantified by Fiji. After imaging, cells were digested with proteinase K (Thermo Fisher Scientific), and the nucleic acids, including the generated biotinylated cDNA, were recovered by phenol-chloroform extraction and concentrated by ethanol precipitation. RNA was digested with RNase H (NEB) and RNase A/T1 (Thermo Fisher Scientific) at 37°C for 1 hour, followed by biotinylated cDNA enrichment using Dynabeads MyOne Streptavidin C1 (Thermo Fisher Scientific). The 3′ cDNA adapter (5′Phos-8 N-AGATCGGAAGAG-CGTCGTGT-3′SpC3) was ligated by T4 RNA ligase 1 (NEB) by incubating at 25°C for 16 hours, and cDNA was recovered with the elution buffer of 95% (v/v) formamide and 10 mM EDTA (pH 8.0) by boiling at 95°C for 10 min, followed by ethanol precipitation. The library was obtained by PCR amplification with next-generation sequencing primers and gel purification of products sized between 180 and 400 bp. Sequencing was performed at the University of Michigan Advanced Genomics Core on a NextSeq platform in single-end mode with 89 bp.

### ARTR-seq Analysis

We adopted the ARTR-seq analytical pipeline^19^ with minor modifications. Raw single-end ARTR-seq FASTQ files were first assessed with FastQC (v0.11.9), and adapter sequences were removed with Cutadapt (v4.3). To deplete ribosomal reads, trimmed reads were aligned to the mouse rRNA reference using STAR (v2.7.10a); reads mapping to rRNA were discarded. The remaining reads were then aligned to the mouse reference genome (GRCm39) with STAR (v2.7.10a), and only alignments with at least 24 matched bases were retained for downstream analysis. Mapped reads were deduplicated by cell barcode and UMI using UMI-tools (v1.1.1), and summarized with featureCounts (v2.0.1). For gene-level analysis, we generated per-gene counts by summing reads mapping to exons and introns of each gene. For intron- and exon-level analyses, we defined canonical exons from Ensembl “canonical” transcripts and treated the intervals between successive canonical exons as introns; featureCounts was used to produce separate count matrices for intronic and exonic regions. Differential enrichment between SRRM2 ARTR-seq libraries and negative primary antibody (IgG) controls was performed with DESeq2 (v1.38.3) using a simple design within each group comparison (0 dpi, 7 dpi, Aged analyzed separately). Features with log2 fold change > 1 and Benjamini–Hochberg–adjusted P value < 0.05 in SRRM2 versus IgG were called nuclear speckle -enriched “DEGs,” and classified as gene-level, intron-level, or exon-level ARTR-seq targets based on the corresponding count table. All other expressed features were considered non–nuclear speckle-enriched and were used as the NS− background in downstream analyses.

### Statistics

Data (mean ± SEM) were analyzed by student’s t-test and two-way ANOVA with Šídák’s multiple comparisons test for parametric distributions, and Mann–Whitney tests for non-parametric distributions. Comparisons of distributions were performed using Wilcoxon rank-sum tests or multiple t-tests with Benjamini–Hochberg correction, with false discovery rate (FDR) or adjusted p values reported as indicated in the figure legends. Significance was set at a p value of 0.05 unless otherwise noted. GraphPad Prism 10 (version 10.1) and R was used for statistical analyses.

## Data Availability

All sequencing data are publicly available at the NCBI Gene Expression Omnibus (GEO) using the accession number GSE317037 and GSE317249. Other data are available upon request.

## Acknowledgments

The authors thank the University of Michigan DNA Sequencing Core for assistance with sequencing library preparation. The authors also thank other members of the Aguilar laboratory for advice and assistance. Research reported in this publication was partially supported by a Hevolution HF-AGE award (C.A.A.), National Science Foundation CAREER award (2045977), Genentech Research Award, the 3M Foundation (C.A.A.), American Federation for Aging Research Grant for Junior Faculty (C.A.A.), the Department of Defense and Congressionally Directed Medical Research Program W81XWH2010336 and W81XWH2110491 (C.A.A.), Defense Advanced Research Projects Agency (DARPA) “BETR” award D20AC0002 (C.A.A.) awarded by the U.S. Department of the Interior (DOI), Interior Business Center, NIH T32 (5T32DE007057) (S.D.G.), NIH T32 (AG000114) (P.D.), and R01HG013495 (C.H.).The content is solely the responsibility of the authors and does not necessarily represent the official views of the National Institutes of Health or National Science Foundation, the position or the policy of the Government, and no official endorsement should be inferred.

## Author Contributions

S.D.G. and P.D. contributed equally to this work. S.D.G., P.D., and C.A.A. conceived and designed the study. S.D.G., P.D., and Y.X. performed experiments. S.D.G., P.D., and Y.S. performed computational and statistical analyses. S.D.G. and P.D. generated figures with input from all authors. S.D.G., P.D., and C.A.A. wrote the first draft of the manuscript, and all authors reviewed, edited, and approved the final version. J.D.W. contributed to study design, data interpretation, and computational analysis strategy. C.H. contributed to study design, ARTR-seq methodology, and data interpretation. C.A.A. supervised the study.

## Competing Interests

C.H. is a scientific founder, a member of the scientific advisory board, and equity holder of AllyRNA Imc., Aferna Bio, Inc. and Ellis Bio Inc., a scientific cofounder and equity holder of Accent Therapeutics, Inc.

