## Supplemental figures for "Nuclear Speckles Regulate Splicing During Muscle Stem Cell Activation and Aging"

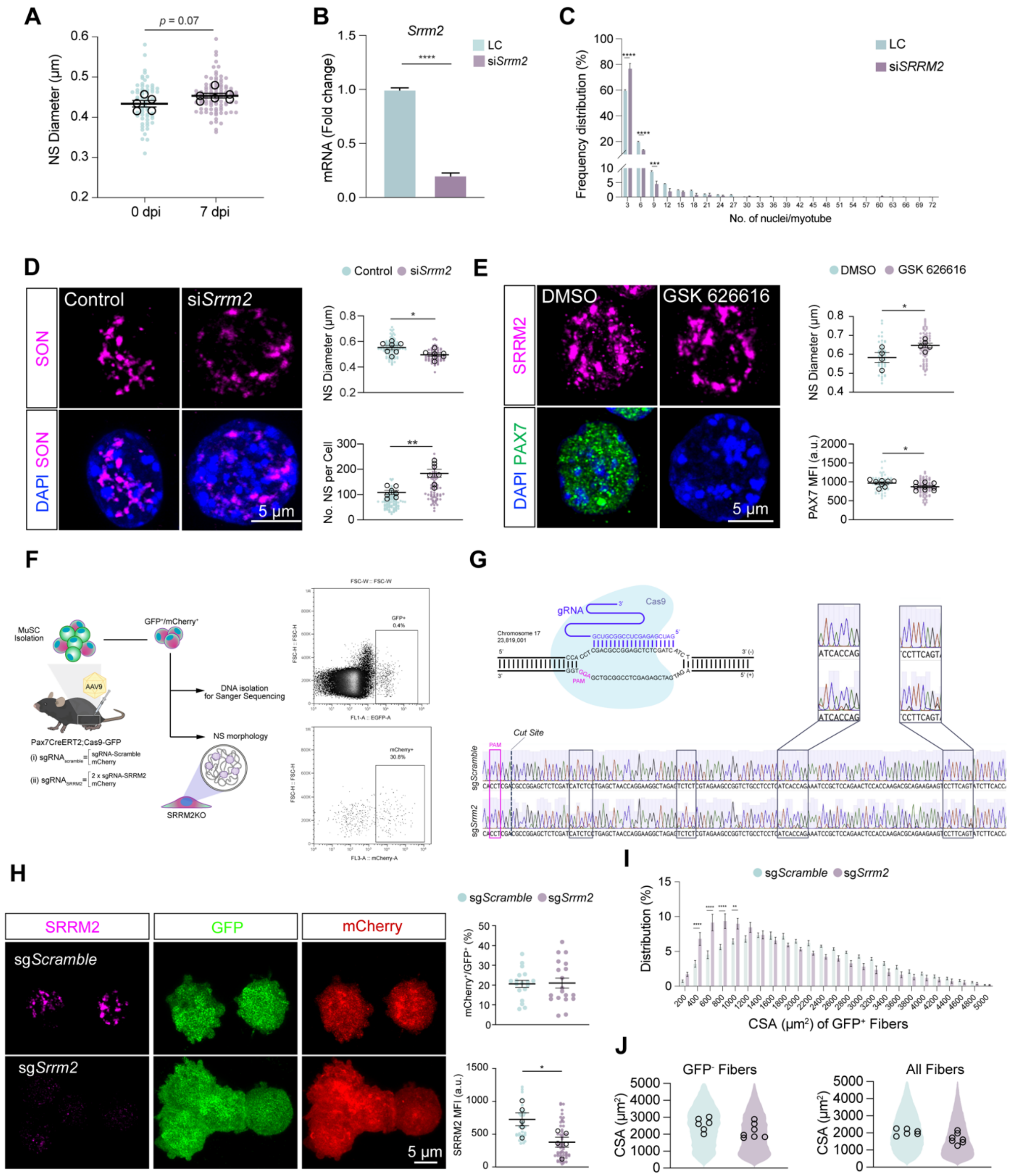

A
p = 0.07
NS Diameter (µm)
0 dpi
7 dpi
B
Srrm2
****
LC
siSrrm2
mRNA (Fold change)
C
LC
siSRRM2
Frequency distribution (%)
No. of nuclei/myotube
D
Control
siSrrm2
SON
DAPI SON
5 µm
NS Diameter (µm)
No. NS per Cell
E
DMSO
GSK 626616
SRRM2
DAPI PAX7
5 µm
NS Diameter (µm)
PAX7 MFI (a.u.)
F
MuSC Isolation
GFP+/mCherry+
AAV9
DNA isolation for Sanger Sequencing
NS morphology
Pax7CreERT2;Cas9-GFP
SRRM2KO
G
gRNA
Cas9
Chromosome 17
PAM
Cut Site
sgScramble
sgSrrm2
H
SRRM2
GFP
mCherry
sgScramble
sgSrrm2
5 µm
mCherry+/GFP+ (%)
SRRM2 MFI (a.u.)
I
sgScramble
sgSrrm2
Distribution (%)
CSA (µm²) of GFP+ Fibers
J
GFP- Fibers
All Fibers
CSA (µm²)

**Figure S1.** (A) Quantification of nuclear speckle diameter between uninjured and injured (7 days post-injury) MuSCs. Single cells= 10-30 per replicate (average of individual NS diameter per cell), replicates= 5-6 animals per group. (B) qPCR of SRRM2 between lipofectamine control and siSRRM2 groups (n= 3 wells per group). (C) Histogram representing the percentage of myotubes with different number of nuclei (field of views= 20-48 per replicate, replicates= 3 animals per group). (D) Single-cell images of NS in lipofectamine control and SRRM2 knockdown (Magenta= SON, Blue= DAPI). Quantification of the diameter (top) and number (bottom) of SON-NS between groups (single cells= 10-18 per replicate, replicates= 7 wells per group). (E) Single-cell images of SRRM2 (top), Pax7 (bottom) between DMSO and GSK 626616. Quantification of the diameter of individual NS (top, single cells= 10-25 per replicate, replicates= 4-5 wells per group) and Pax7 MFI (bottom, field of views= 6-20 per replicate, replicates= 8-9 wells per group). (F) AAVs with a gRNA targeting SRRM2 or a non-targeting/scramble gRNA were injected intramuscularly to the tibialis anterior and gastronecmius muscles to Pax7-Cas9 mice. Three weeks after injections, transduced MuSCs were isolated via FACS to assess edit with sanger sequencing and decrease in protein expression with immunocytochemistry. Example of gating strategy to isolate $GFP^+$ (top) and $mCherry^+/GFP^+$ cells. (G) Schematic describing the CRISPR-Cas9 edit (top). Examples of traces from scramble (top) and SRRM2-KO (bottom) groups for one of the gRNAs (GATCGAGAGCTCCGGCGTCG). Dash line represents the Cas9 cut site (bottom). (H) Representative images of $mCherry^+/GFP^+$ MuSCs after delivery of sgScramble (top) or sgSRRM2 (bottom). Quantification of the percentage of $mCherry^+$ cells (top, n=18-19 animals per group) and mean fluorescence intensity of SRRM2 between groups (bottom, single cells= 8-12 per replicate, replicates= 5 animals per group). (I) Histograms of $GFP^+$ fibers representing percentages of fibers with the corresponding fiber areas. (J) Quantification of individual cross-sectional area of $GFP^-$ only fibers and all fibers ($GFP^+$ and $GFP^-$) across groups. Violin plot represents the overall distribution of all replicates between scramble and SRRM2-KO groups and circles indicate the median of each replicate (n= 6-7 animals per group). *p*-values derived from student's t-test and two-way ANOVA with Šídák's multiple comparisons test. **p*<0.05, ***p*<0.01, *****p*<0.0001; mean ± SEM.

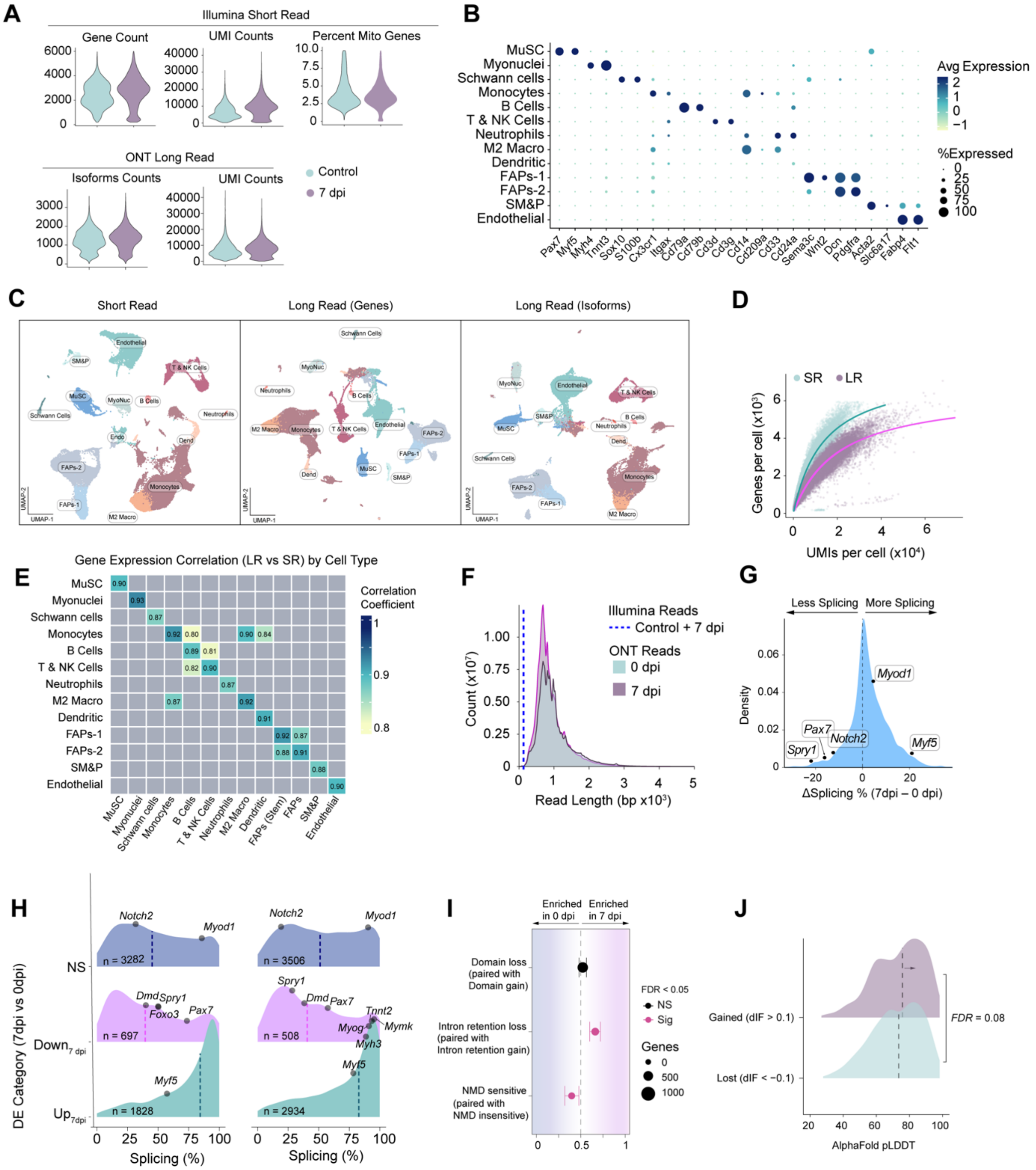

A
Illumina Short Read
Gene Count
UMI Counts
Percent Mito Genes
ONT Long Read
Isoforms Counts
UMI Counts
Control
7 dpi
B
MuSC
Myonuclei
Schwann cells
Monocytes
B Cells
T & NK Cells
Neutrophils
M2 Macro
Dendritic
FAPs-1
FAPs-2
SM&P
Endothelial
Avg Expression
%Expressed
Pax7 Myf5 Myh4 Tnnt3 Sox10 S100b Cx3cr1 Itgax Cd79a Cd79b Cd3d Cd3g Cd14 Cd209a Cd33 Cd24a Sema3c Wnt2 Dcn Pdgfra Acta2 Slc6a17 Fabp4 Flt1
C
Short Read
Long Read (Genes)
Long Read (Isoforms)
D
SR
LR
Genes per cell (x10³)
UMIs per cell (x10⁴)
E
Gene Expression Correlation (LR vs SR) by Cell Type
Correlation Coefficient
F
Illumina Reads
Control + 7 dpi
ONT Reads
0 dpi
7 dpi
Count (x10⁷)
Read Length (bp x10³)
G
Less Splicing
More Splicing
Density
ΔSplicing % (7dpi – 0 dpi)
Myod1
Pax7
Notch2
Spry1
Myf5
H
DE Category (7dpi vs 0dpi)
NS
Down7 dpi
Up7dpi
n = 3282
n = 697
n = 1828
n = 3506
n = 508
n = 2934
Splicing (%)
I
Enriched in 0 dpi
Enriched in 7 dpi
Domain loss (paired with Domain gain)
Intron retention loss (paired with Intron retention gain)
NMD sensitive (paired with NMD insensitive)
FDR < 0.05
NS
Sig
Genes
J
Gained (dIF > 0.1)
Lost (dIF < −0.1)
FDR = 0.08
AlphaFold pLDDT

**Figure S2.** (A) Violin plots showing short-read quality control metrics (gene count, UMI counts, and percent mitochondrial genes) and long-read quality control metrics (isoform counts and UMI counts) across 0 dpi and 7 dpi samples. (B) Dot plot of canonical marker gene expression across annotated cell types in the integrated dataset; color indicates average expression and dot size indicates percentage of cells expressing each gene. (C) UMAPs of short-read gene expression, long-read gene expression, and long-read isoform expression embeddings, each colored by annotated cell type. (D) Curves showing the relationship between genes detected per cell and UMI counts per cell for short-read and long-read data. (E) Heatmap of Pearson correlations between short-read and long-read gene expression profiles across annotated cell types. (F) Read length distributions for Illumina short reads and Oxford Nanopore long reads across 0 dpi and 7 dpi samples. (G) Density plot of ΔSplicing% (7 dpi − 0 dpi) for genes detected in both conditions, with representative quiescence and activation genes labeled. (H) Density plots of junction-weighted splicing percentage stratified by differential gene expression category between 7 dpi and 0 dpi, with non-significant genes (NS), genes downregulated at 7 dpi, and genes upregulated at 7 dpi shown separately. Left, all genes with long-read coverage; right, genes shared between conditions. (I) Dot plot summarizing significant isoform switches grouped by functional consequence, including domain loss paired with domain gain, intron retention loss paired with intron retention gain, and NMD-sensitive paired with NMD-insensitive isoforms; x axis indicates enrichment in 0 dpi or 7 dpi, point size reflects the number of genes, and color denotes significance. (J) Density plot of AlphaFold pLDDT values for isoforms that gained usage (dIF > 0.1) or lost usage (dIF < -0.1) between 0 dpi and 7 dpi.

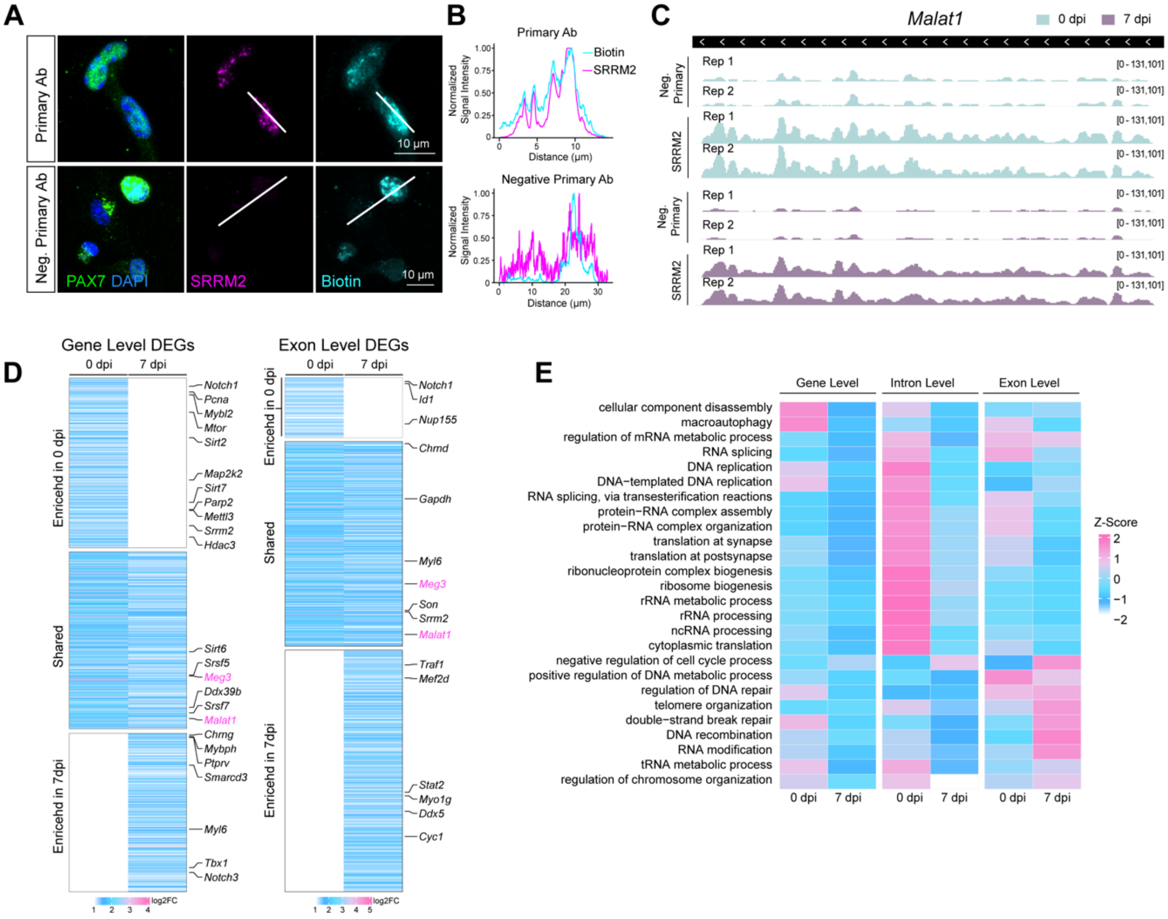
A
Primary Ab
Neg. Primary Ab
PAX7 DAPI
SRRM2
Biotin
10 μm
B
Primary Ab
Biotin
SRRM2
Normalized Signal Intensity
Distance (μm)
Negative Primary Ab
C
Malat1
0 dpi
7 dpi
Neg. Primary
SRRM2
Rep 1
Rep 2
[0 - 131,101]
D
Gene Level DEGs
Exon Level DEGs
0 dpi
7 dpi
Enricehd in 0 dpi
Shared
Enricehd in 7dpi
Notch1
Pcna
Mybl2
Mtor
Sirt2
Map2k2
Sirt7
Parp2
Mettl3
Srrm2
Hdac3
Sirt6
Srsf5
Meg3
Ddx39b
Srsf7
Malat1
Chrng
Mybph
Ptprv
Smarcd3
Myl6
Tbx1
Notch3
Id1
Nup155
Chrnd
Gapdh
Son
Traf1
Mef2d
Stat2
Myo1g
Ddx5
Cyc1
log2FC
E
Gene Level
Intron Level
Exon Level
cellular component disassembly
macroautophagy
regulation of mRNA metabolic process
RNA splicing
DNA replication
DNA−templated DNA replication
RNA splicing, via transesterification reactions
protein−RNA complex assembly
protein−RNA complex organization
translation at synapse
translation at postsynapse
ribonucleoprotein complex biogenesis
ribosome biogenesis
rRNA metabolic process
rRNA processing
ncRNA processing
cytoplasmic translation
negative regulation of cell cycle process
positive regulation of DNA metabolic process
regulation of DNA repair
telomere organization
double−strand break repair
DNA recombination
RNA modification
tRNA metabolic process
regulation of chromosome organization
Z-Score

**Figure S3.** (A) Representative images showing biotin signal, SRRM2 immunostaining, and line-scan quantification in MuSCs processed with SRRM2 primary antibody or negative primary antibody control. (B) Line-scan plots of normalized biotin and SRRM2 signal intensity for primary antibody and negative primary antibody control conditions. (C) Genome browser views of *Malat1* ARTR-Seq signal in 0 dpi and 7 dpi samples for negative primary antibody control and SRRM2 pulldown, shown across biological replicates. (D) Heatmaps of gene-level differentially enriched genes (left) and exon-level differentially enriched genes (right) from ARTR-Seq, grouped by enrichment at 0 dpi, shared across 0 and 7 dpi, or enriched at 7 dpi, with selected genes labeled. (E) Heatmap of gene ontology enrichment z scores for gene-level, intron-level, and exon-level ARTR-Seq datasets at 0 dpi and 7 dpi.

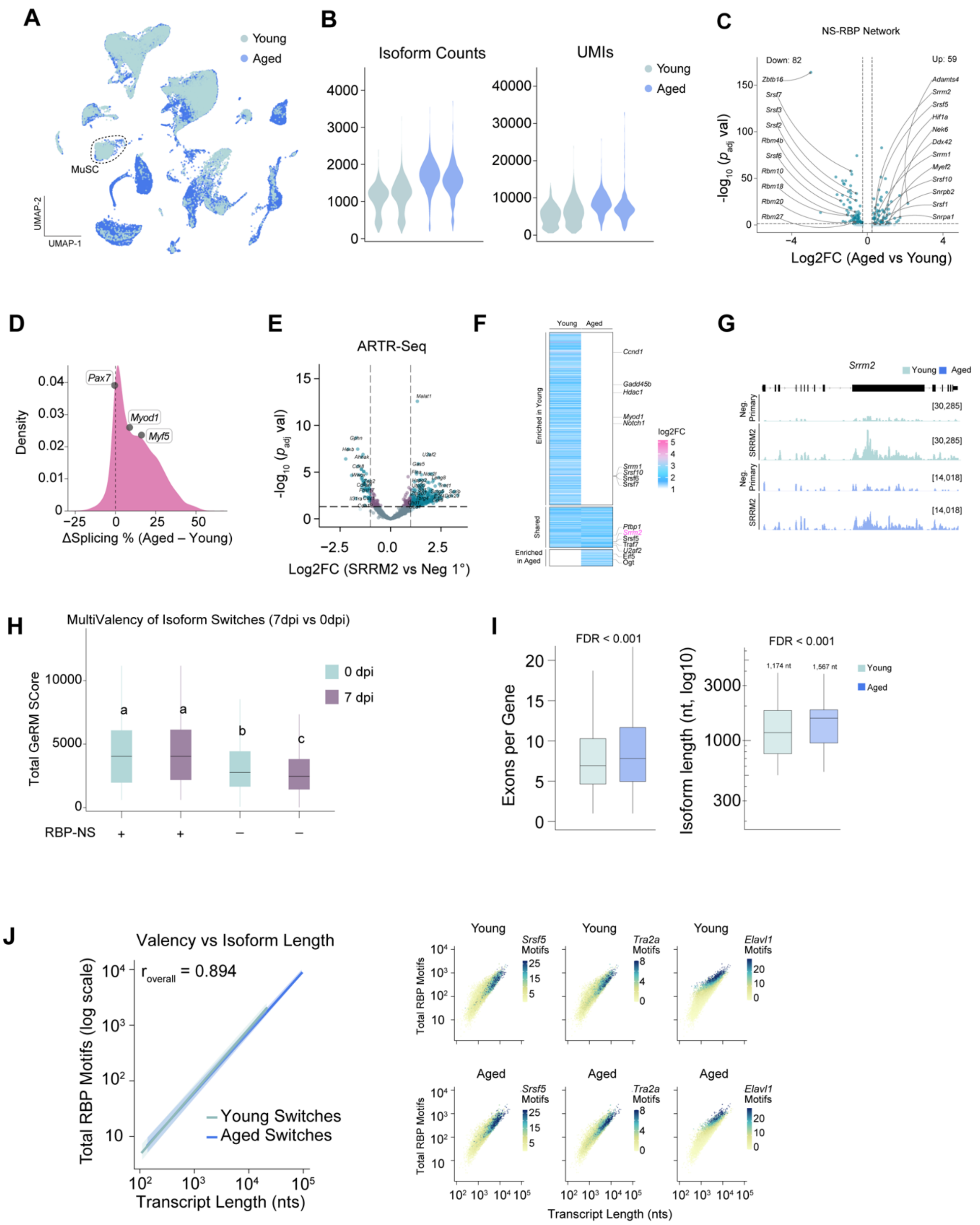
A
Young
Aged
MuSC
UMAP-2
UMAP-1
B
Isoform Counts
UMIs
Young
Aged
C
NS-RBP Network
Down: 82
Up: 59
-log10 (p adj val)
Log2FC (Aged vs Young)
D
Pax7
Myod1
Myf5
Density
ΔSplicing % (Aged – Young)
E
ARTR-Seq
-log10 (p adj val)
Log2FC (SRRM2 vs Neg 1°)
F
Young
Aged
Enriched in Young
Shared
Enriched in Aged
log2FC
G
Srrm2
Young
Aged
Neg. Primary
SRRM2
H
MultiValency of Isoform Switches (7dpi vs 0dpi)
Total GeRM SCore
0 dpi
7 dpi
RBP-NS
I
FDR < 0.001
Exons per Gene
Isoform length (nt, log10)
1,174 nt
1,567 nt
Young
Aged
J
Valency vs Isoform Length
r overall = 0.894
Total RBP Motifs (log scale)
Transcript Length (nts)
Young Switches
Aged Switches
Young
Aged
Srsf5 Motifs
Tra2a Motifs
Elavl1 Motifs
Total RBP Motifs
Transcript Length (nts)

**Figure S4.** (A) UMAP of integrated Young and Aged single-cell data colored by age group, with MuSCs indicated. (B) Violin plots showing long-read isoform counts and UMI counts in young and aged samples. (C) Volcano plot of differential expression within the NS-RBP network comparing Aged versus Young MuSCs, with selected genes labeled. (D) Density plot of ΔSplicing% (Aged − Young) for genes detected in both conditions, with representative genes labeled. (E) Volcano plot of SRRM2 ARTR-Seq enrichment over negative primary antibody background in aged MuSCs, with selected genes labeled. (F) Heatmap of intron-level differentially enriched genes from ARTR-Seq in young and aged MuSCs, grouped by enrichment in young, shared, or enriched in aged, with selected genes labeled. (G) Boxplots of total generalized RNA multivalency (GeRM) scores for isoform switches in the 0 dpi versus 7 dpi comparison, stratified by RBP–NS membership. (H) Boxplots comparing exon number per gene and isoform length for switched isoforms in young and aged MuSCs. (I) Relationship between transcript length and total RNA binding protein motif abundance across isoform switches, shown for Young and Aged MuSCs overall (left) and for representative motifs (*Srsf5*, *Tra2a*, and *Elavl1*) in each condition (right).
